# The NQR complex regulates the immunomodulatory function of *Bacteroides thetaiotaomicron*

**DOI:** 10.1101/2022.10.21.513251

**Authors:** Morgan J. Engelhart, Robert W. P. Glowacki, Jessica M. Till, Clifford V. Harding, Eric C. Martens, Philip P. Ahern

**Affiliations:** Department of Cardiovascular and Metabolic Sciences, Lerner Research Institute, Cleveland Clinic, Cleveland, OH 44195, USA; Department of Molecular Medicine, Lerner College of Medicine, Case Western Reserve University, Cleveland, OH; Center for Microbiome and Human Health, Lerner Research Institute, Cleveland Clinic, Cleveland, OH 44195, USA; Department of Pathology, Case Western Reserve University/University Hospitals Cleveland Medical Center, Cleveland, Ohio, USA; Department of Microbiology and Immunology, University of Michigan Medical School, Ann Arbor, MI 48109, USA

## Abstract

The gut microbiome and intestinal immune system are engaged in a dynamic interplay that provides myriad benefits to host health. However, the microbiome can also elicit damaging inflammatory responses, and thus establishing harmonious immune-microbiome interactions is essential to maintain homeostasis. Gut microbes actively coordinate the induction of anti-inflammatory responses that establish these mutualistic interactions. Despite this, the microbial pathways that govern this dialogue remain poorly understood. We investigated the mechanisms through which the gut symbiont *Bacteroides thetaiotaomicron* exerts its immunomodulatory functions. Our data reveal that *B. thetaiotaomicron* stimulates production of the cytokine IL-10 via secreted factors that are packaged into outer membrane vesicles, in a TLR2 and MyD88 dependent manner. Using a transposon mutagenesis based screen, we identified a key role for the *B. thetaiotaomicron* encoded NQR complex, which regenerates NAD+ during respiration, in this process. Finally, we found that disruption of NQR reduces the capacity of *B. thetaiotaomicron* to induce IL-10 by impairing biogenesis of outer membrane vesicles. These data identify a microbial pathway with a previously unappreciated role in gut microbe mediated immunomodulation that may be targeted to manipulate the capacity of the microbiome to shape host immunity.

**Key points:** The *B. theta* NQR complex coordinates OMV-driven TLR2-dependent IL-10 expression.

## Introduction

The gastrointestinal tract is densely colonized with a complex array of bacteria, fungi, archaea, and viruses collectively referred to as the gut microbiome (1–3). The intestine is also home to a multifaceted immune network that is separated from the gut microbiome by an overlying epithelial and mucus layer (4, 5). Despite this physical barrier, the intestinal immune system is in continual contact with the microbiome. Given the important contribution of the microbiome to maximizing host health, it is essential that the host maintains harmonious relations with these microbial communities to sustain a mutualistic state, while remaining vigilant for potential infectious agents. To this end, the intestinal immune system is enriched with cell types and molecules that prevent unwarranted deleterious inflammatory responses, and limit the penetration of the microbiome into host tissues (6). This includes a large population of CD4^+^ regulatory T cells (Tregs) (7–14), and expression of cytokines such as IL-10 (13, 15) and transforming growth factor-beta (TGF-β) (16), potent anti-inflammatory mediators. The essential nature of these responses is highlighted by the deleterious impact of defects in these pathways, which leads to the development of tissue-destructive, microbiome-targeted pro-inflammatory immune responses, as occurs in inflammatory bowel disease (17–27). However, the molecular factors that drive the accumulation/production of these tolerance inducing factors, and the microbial pathways involved, remain poorly understood.

A large body of evidence has established that specific gut microbes actively coordinate the activation of host anti-inflammatory responses. These include, bacteria such as *Helicobacter hepaticus* (15, 28, 29), *Bacteroides fragilis* (13, 30–33), *Bacteroides thetaiotaomicron* (*B. theta* hereafter) (9, 34, 35), *Bifidobacterium bifidum* (14), members of the altered Schaedler flora (ASF) (10) and collections of *Clostridium* species (7, 8, 36) among others, that have been shown to induce the accumulation of colonic Tregs and/or production of the anti-inflammatory cytokine IL-10. Immunomodulatory microbiome-derived products identified to date have fallen into two main categories (i) products of microbial metabolism like short-chain fatty acids or microbial-modified bile acids (36–43) and (ii) cellular components like capsular or cell-surface polysaccharides (13-15, 31, 32) that are sensed by host pattern recognition receptors (PRRs) (14, 15, 32, 33, 44). Although our knowledge of the precise pathways through which the host is exposed to these molecules is limited, outer membrane vesicles (OMVs), spherical structures derived from the outer membrane of many Gram-negative bacteria (33, 45–49), are posited to be packaged with proteins, lipids, and polysaccharides, and delivered to the intestinal immune system to shape the local immunological milieu. Indeed, members of the *Bacteroides*, a predominant genera within the gut microbiome (50–52), including *B. fragilis* (33) and *B. theta* (45, 53), produce OMVs that contain immunomodulatory factors that can invoke beneficial immune response that limit inflammation, *e.g.* OMVs from *B. fragilis* contain polysaccharide A (PSA) that elicits IL-10 production (33).

Given their enormous potential to shape immune responses, gut microbes and/or their derived products have the potential to act as novel therapeutics for inflammatory diseases by favoring the induction of anti-inflammatory responses. Despite this promise, and notwithstanding some key advances, our understanding of these molecules, and how they are delivered to the host, remains limited. As our knowledge of the host-microbiome dialogue that promotes mutualism grows, there is an increasing need to understand the microbial pathways that govern the production and secretion of these factors. Defining such pathways will provide an opportunity to potentially expand production of molecules of interest, or heterologously reconstruct these pathways in synthetic probiotics to maximize beneficial effects (54).

We have leveraged the genetically tractable gut bacterium, *B. theta*, a prominent microbiome member that we and others (9, 34, 35, 41) have shown to induce anti-inflammatory responses, to understand the microbial pathways that impact gut bacterial interaction with the immune system. Our data have identified that *B. theta* induces the anti-inflammatory cytokine IL-10 via secreted factors in a Toll-like receptor 2 (TLR2)-MyD88 dependent manner. In addition, we have identified that the Na^+^-translocating <u>N</u>ADH:ubi<u>q</u>uinone oxidoreductase (*nqr*) locus (55–57)—encoded by genes *BT1155-BT1160*—is required for the optimal induction of IL-10. We show that disruption of the *nqr* operon impairs OMV biogenesis and that this altered OMV generation is responsible for the reduced IL-10 induction. Disruption of the NQR complex indirectly alters OMV biogenesis to regulate IL-10 induction, rather than through a direct NQR associated mechanism such as NAD+ regeneration. Our results provide a formal demonstration that couples OMV production with the induction of IL-10 by members of the microbiome, and offer new insights into the pathways that regulate the generation of OMVs. Thus, the NQR complex represents a potential microbiome target that could alter host-microbe interactions and favor the promotion of an anti-inflammatory intestinal environment.

## Methods

### Bacterial strains, culturing conditions, and molecular genetics

*Bacteroides thetaiotaomicron* ATCC 29148 (VPI-5482) and its genetic variants were grown in tryptone-yeast extract-glucose (TYG) broth medium or on brain heart infusion (BHI) agar (BD, 211059) supplemented with 10% v/v defibrinated horse blood (Quad Five, Cat. # 210-500) at 37°C in an anaerobic chamber (Compressed gas: 5% CO^2^, 5% H_2_, 90% N_2_, chamber set to ∼2.5% H_2_, Coy Manufacturing, Grass Lake, MI). TYG was made using: Tryptone Peptone (10 g/L; Gibco, Cat. # 211921), Bacto Yeast Extract (5 g/L; BD, Cat. # 212750), Glucose (2 g/L; Sigma, Cat. # G8270), Cysteine HCl (0.5 g/L; Sigma, Cat. # C1276), 1M Potassium Phosphate Buffer (pH 7.2, 100 mL/L) made using 1M potassium phosphate monobasic (Sigma, Cat. # P0662), and 1M potassium phosphate dibasic (Sigma, Cat. # P3786), Vitamin K_3_ (menadione) (1 mL/L of a 1 mg/mL stock which was prepared in 100% ethanol; Sigma, Cat. # M5625), TYG Salts (40 mL/L) made using MgSO_4_.7H_2_O (0.5 g/L; Sigma, Cat. # M63-500), NaHCO_3_ (10 g/L; Sigma, Cat. # S4019), and NaCl (2 g/L; Sigma, Cat. # S3014), FeSO_4_ (1 mL/L of a 0.4 mg/mL stock; Sigma, Cat. # F8048), CaCl_2_ (1 mL/L of an 8 mg/mL stock; Sigma, Cat. # C7902), Hemin (0.5 mL/L of a 10 mg/mL stock which was prepared in 0.1 N NaOH; Sigma, Cat. # 51280). All solutions were made in Milli-Q-water unless otherwise noted. Genetic deletions were performed by counter-selectable allelic exchange as previously described (58), and all bacterial strains, plasmids, and primers are referenced in Supplemental Table 1. Briefly, deletion of genes encoding the NQR complex was done using a modified form of *B. theta* strain VPI-5482 with a deletion of the thymidine kinase gene *tdk* (*BT2275*) to facilitate counter-selection. In addition, deletion of genes encoding the NQR complex were also made as indicated using *B. theta*^Δ*tdk*Δ*cps*1-8^ as the parent strain (59). Deletion of genes or bacterial gene clusters *BT1160, BT1161, BT1155-BT1159,* and *BT1155-BT1160,* was completed by ligating PCR-amplified 750bp fragments flanking the gene of interest into the suicide vector (pExchange-*tdk*) using Sal1 (NEB, Cat. # R0138L) and Xbal (NEB, Cat. # R0145L) cut sites. This vector contains a cloned copy of *tdk* (provides resistance to the toxic nucleotide analog 5-fluoro-2’-deoxyuridine (FUdR))*, bla* (ampicillin resistance), and *ermGb* (erythromycin resistance), to facilitate selection and counter-selection. The resulting construct was conjugated into either *B. theta*^Δ*tdk*^ or *B. theta*^Δ*tdk*Δ*cps*1-8^. Individual-recombinant merodiploid colonies, selected on erythromycin (25 μg/mL), were pooled and plated on BHI-blood agar containing FUdR (200 μg/mL) to select for recombinants. Candidate gene deletions were screened by PCR and DNA sequencing to identify isolates that had the desired gene deletions.

Complementation of gene *BT1160* was accomplished as previously described in (60). Briefly, wild-type alleles encoding *BT1160* and promoter regions were amplified using the primers listed in Supplemental Table 1. Complementing alleles were ligated into the *pNBU2*-*ermGb* vector using Sal1 and BamH1 (NEB, Cat. # R0136L) cut sites and the resulting construct was inserted into *att2* (tRNA^ser^) in *B. theta*^Δ*tdk*Δ*BT1160_*trunc^. Individual recombinant colonies, selected on erythromycin (25 μg/mL) were screened by PCR and DNA sequencing to identify successfully complemented isolates.

### *B. theta* heat-inactivated pellets and conditioned media

*B. theta* was grown anaerobically in TYG growth medium overnight to stationary phase (∼16 hours) and the OD_600_ measured (ranging from ∼2.0-3.5). When comparing the effects of multiple *B. theta* strains or mutants on cytokine induction, cultures were normalized by OD using TYG. Normalized cultures were spun down at 7000 *x g* for 5 minutes at 4°C, and the supernatant was separated from the pellet by pouring off into a fresh tube. The supernatant was filtered through a 0.22 μm PVDF filter (Millex, Cat. # SLGVR33RS) to make aliquots of cell-free sterile supernatant, referred to as conditioned media hereafter. Conditioned media was frozen at -20°C until further use. The bacterial pellet was washed by resuspending in sterile PBS, spun down as before, and resuspended in a volume of sterile PBS equivalent to the volume of media from which the pellet was obtained, the OD_600_ was measured again, and additional dilutions were made in PBS such that equivalent CFU (based on OD_600_) of different strains or mutants were used to stimulate cells in downstream assays. Aliquots of the washed and OD_600_ normalized pellets (1 mL) were transferred to sterile Eppendorf tubes and placed at 70°C for 30 minutes to heat-inactivate them. Heat-inactivated pellets were stored at -20°C until use. The effectiveness of sterile filtering the conditioned media was confirmed by plating on BHI blood agar plates and incubated at 37°C anaerobically for >3 days (data not shown).

### Outer Membrane Vesicle (OMV) isolation

OMVs were isolated from conditioned media using an ultracentrifugation based approach as previously described (61). A set volume of conditioned media was placed into 1.5 mL ultracentrifugation tubes (ThermoScientific, Cat. # 314352H01) and spun at 100,000 *x g* for 2 hours at 4°C. Following the spin, the supernatant was removed and the pellet (containing the OMVs) was resuspended and washed in 1 mL of sterile PBS (all PBS was first double filtered through a 0.1 μm PES filter (Thermo Scientific; Cat. #5670010) before use). This resuspension was spun down again at 100,000 *x g* for 2 hours, the supernatant was removed, and pellet containing the washed OMVs was resuspended in 0.1 μm filtered PBS. OMVs were either concentrated (for Qubit concentration analysis, see below) or resuspended in a volume of PBS equivalent to the volume of conditioned media from which they were extracted (for IL-10 induction assay, see below). Isolated OMVs were stored at -20°C until further use.

### *In vitro* IL-10 assay

The ability of the heat-inactivated pellets and conditioned media from *B. theta* strains or mutants to induce IL-10 was tested with an *in vitro* assay using bulk unfractionated murine splenocytes. Splenocytes were isolated from the spleen of wild-type, TLR2^-/-^, TLR4^-/-^, MyD88^-/-^, or Dectin-1^-/-^ mice by forcing through a 70 μm cell-strainer using sterile PBS containing 0.1% w/v Bovine Serum Albumin (BSA; Sigma). Cells were pipetted up and down to generate a single cell suspension and collected in a 15 mL conical tube, and pelleted by centrifugation at 454 *x g* for 5 minutes. The supernatant was removed, the pellet was resuspended by flicking the tube and 750 μL (per spleen) of room temperature ACK lysis buffer (Gibco, Cat. # A10492) was added, followed by immediate vortexing. The cells were incubated for 3 minutes at room temperature, and ∼10 mL of PBS/0.1% BSA was then added to the cells to halt lysis. The cell suspension was then passed through a 70 μm cell-strainer and the flow-through collected into a new 15 mL conical tube. Cells were pelleted as before, the supernatant removed, and the pellet was resuspended in 5 mL complete RPMI (RPMI 1640 supplemented with L-glutamine, 20 mM final concentration HEPES (Gibco, Cat # 15630-080), 10% v/v final concentration heat-inactivated fetal bovine serum (FBS), 100 units/mL Penicillin and 100 μg/mL Streptomycin) per spleen. Live cells were counted by hemocytometer (stained with trypan blue) and diluted with complete RPMI to 5x10^6^ cells/mL.

Splenocytes were plated in a flat-bottomed sterile 96-well tissue culture treated plate (Gibco, Cat. # 353072) by adding 100 μL to each well for a final concentration of 5x10^5^ cells/well. Splenocytes were then incubated with heat-inactivated bacterial pellets (OD_600_ 0.05-1.0), conditioned media (0.1-7% v/v), or OMVs (0.1-5% v/v) and complete RPMI was added to each well to bring the final volume to 200 μL. Cells and samples were incubated at 37°C in 5% CO_2_ for 48 hours, following which the cells were pelleted by centrifugation at 454 *x g* for 5 minutes and the supernatants collected. Supernatants were frozen at -20°C until further processing, and IL-10 was measured by ELISA (Biolegend, Cat. # 431411).

The *in vitro* IL-10 assay was also performed using bone-marrow derived macrophages (BMDMs) from wild-type or TLR2^-/-^ mice. Briefly, bone marrow was isolated from the femur and tibia using a tube-in-tube centrifugation method, where a hole was poked in the bottom of a sterile PCR tube using a sterile 18G needle which was then placed in a sterile Eppendorf tube. The ends of the femur and tibia were removed and the bones were placed open side down in the PCR tube. The tubes were centrifuged briefly to remove the bone marrow and lysed with ACK lysis buffer (as described above; 100 μL per tube). Lysis was halted with PBS/0.1% BSA and all the bone marrow was combined, cells were passed through a 70 μm cell-strainer and the flow-through was collected into a new 15 mL conical tube. Cells were pelleted at 454 *x g* for 5 minutes before being resuspended in BMDM media (DMEM-low glucose with L-glutamine and sodium pyruvate, 20% v/v FBS, 100 units/mL Penicillin, 100 μg/mL Streptomycin, 30% v/v L929-cell media [macrophage colony-stimulating factor (M-CSF) is secreted by the L929-cell media (LCM) (62) to support BMDM development], 20 mM HEPES). LCM was generated by harvesting and sterile filtering the supernatant from confluent cell culturing of L929 cells (NCPC L929, ATCC # CCL-1) in DMEM media (DMEM-low glucose with L-glutamine and sodium pyruvate, 100 units/mL Penicillin and 100 μg/mL Streptomycin, 10% v/v FBS). ACK lysed bone marrow was plated at 2x10^6^ cells per non-tissue culture treated petri dish in 10 mL BMDM media. The media was changed every three days, and the macrophages were harvested using versene (Gibco, Cat. #15040-066) after day 10 when the cells were adherent and elongated. BMDMs were plated at a final concentration of 5x10^5^ cells/well, and the supernatant was collected after 2 days for detection of IL-10. All spleens/bone marrow were isolated from animals whose use was approved by the Institutional Animal Care and Use Committee (IACUC) of the Cleveland Clinic (wild-type [Jackson Labs, strain #: 000664], TLR4-/- [Jackson Labs, strain #: 029015](63), and TLR2-/- [Jackson Labs, strain #: 004650](64)), the IACUC of Case Western Reserve University (TLR2-/- and MyD88-/- [generously provided to C.V.H. by Drs. O. Takeuchi and S. Akira, Research Institute for Microbial Disease, Osaka University, Osaka, Japan, and back-crossed to C57BL/6J mice a minimum of eight times as per (63, 65–67)]) or the IACUC of the University of Minnesota (Jackson Labs, wild-type [strain #: 000664] and Dectin-1-/- [Jackson Labs, strain #: 012337](68)). Dectin-1-/- mice are reported by Jackson Labs to be on a mixed C57BL/6N;C57BL/6J genetic background. All mice used were male unless otherwise indicated.

*In vitro* IL-10 assays were also performed using human Peripheral Blood Mononuclear Cells (PBMCs) (STEMCELL Technologies, Cat. # 70025.1). The assay was performed as described above, and PBMCs were thawed and handled according to the manufacturer instructions. Briefly, the outside of the frozen vial of cells was wiped with 70% ethanol, vented in a biosafety cabinet by quickly uncapping the tube to release built up pressure, and then recapped. The cells were thawed at 37°C in a water bath by gently shaking the vial. The vial was removed from the water bath while a small frozen cell pellet remained. The outside of the vial was cleaned with 70% ethanol again. Cells were transferred to a 50 mL conical and 1 mL of complete RPMI was added to the cells dropwise. Cells were washed by slowly adding 15-20 mL complete RPMI dropwise while swirling the tube. Cells were spun down at 454 *x g* for 10 minutes at room temperature and the supernatant was carefully removed. Cells were resuspended by gently flicking the tube in the remaining volume. 100 μg/mL of DNase 1 (STEMCELL Technologies, Cat. # 07900) was added to the cell suspension for 15 minutes at room temperature to prevent cell clumping. Cells were washed with 15-20 mL complete RPMI and spun as described above. The supernatant was removed and the pellet was resuspended by gently flicking the tube. The volume was measured and live cells were counted by hemocytometer (stained with trypan blue) and diluted with complete RPMI to 5x10^6^ cells/mL. PBMCs were then plated as described above and human IL-10 was measured in the cell supernatants by ELISA (Biolegend, Cat. # 430604).

### IL-10 induction by transposon mutagenesis library

The ability of *B. theta* transposon mutants to induce IL-10 was tested using an arrayed *B. theta*^Δ*tdk*Δ*cps*1-8^ transposon mutagenesis library. The library was previously generated using the *mariner* transposon (35). Each 96-well plate of mutants was grown anaerobically at 37°C in TYG media, to stationary phase, overnight, and frozen at -80°C until use. The plates were thawed and spun at 454 *x g* for 5 minutes to separate the bacterial pellet from the TYG media supernatant. Splenocytes were stimulated as described above using a dose of 1% v/v conditioned media or Pam3CSK4, a defined TLR2 agonist (69–71) (each plate had wells stimulated with Pam3CSK4 as a positive control), and IL-10 induction was measured in the supernatant after 2 days by ELISA. Each mutant was tested in duplicate, and we then assessed IL-10 induction in culture supernatants by ELISA, and normalized the values for each transposon mutant to the positive control, Pam3CSK4, to account for plate-to-plate variation. In addition, we performed growth assessment of the individual mutants and 27 mutants that did not exhibit detectable growth were removed from our analysis. The data was processed with custom R script (https://github.com/PhilipAhern/Engelhart-et-al.git). Briefly, the normalized IL-10 expression levels were converted to log2 scale for normalization, and then any transposon mutant with an IL-10/Pam3CSK4 ratio ≥2 standard deviations below the mean were filtered, and z-score and p-value were computed for each using normal distribution statistics. Adjusted p-values were calculated by the false discovery rate (FDR) to correct for multiple hypothesis testing. Transposon mutant 19_H4 was sent for whole genome sequencing (Illumina; 2 X 151 bp) to identify the insertion site of the transposon.

### *Nqr* locus gene expression

RNA was isolated from three 5 mL cultures of *B. theta*^Δ*tdk*Δ*BT1160_*trunc^, *B. theta*^Δ*tdk*Δ*BT1160*_full^, and their parent strains (*B. theta*^Δ*tdk*^) grown to mid-log phase (OD 0.5-0.8) in TYG. The 5 mL cultures were transferred to a 15 mL conical tube and spun down at 7000 *x g* for 5 minutes at room temperature. The supernatant was removed (note, a 1 mL aliquot was sterile filtered and kept to test conditioned media for IL-10 induction (as above)) and replaced with 5 mL of RNA-protect (Qiagen, Cat. # 1018390), mixed by pipetting, and allowed to sit for 5 minutes at room temperature. Samples were spun for 10 minutes at 7000 *x g* at room temperature and the supernatant was decanted, and all additional liquid was removed by gently taping the tube on a paper towel. Samples were stored at -80°C until RNA was isolated.

RNA was isolated following manufactures instructions provided by RNeasy kit (Qiagen, Cat. # 74004). Briefly, pellets were brought to room temperature and resuspended in 200 μL of TE buffer containing 1 mg/mL lysozyme (Sigma, Cat. #L4919) by pipetting, and allowed to incubate for 5 minutes at room temperature, pipetting for 10 seconds every 2 minutes. 700 μL of RLT buffer (containing β-mercaptoethanol (BME) 10 μL/mL of a 14.3 M stock solution) was added to the solution and mixed by pipetting, followed by 500 μL of 100% ethanol and mixed by pipetting. The solution was transferred to an RNeasy Mini Spin column (Econospin Cat. # 1940-250) and spun for 15 seconds at 18,111 *x g* and the flow-through was discarded. 700 μL of Buffer RW1 was added to the column and spun again, as above, to wash the membrane. The flow through was discarded and 500 μL of Buffer RPE was added to the column and spun again. The flow-through was discard and this step was repeated followed by a 2 minute spin. The flow-through was discarded and the tube was spun again for 1 minute to remove any residual ethanol. The column was placed in an RNase-free 1.5 mL collection tube (Ambion, Cat. # AM12400) and 30 μL of nuclease free water was added to the column membrane, let to sit for 5 minutes at room temperature and then spun at 8476 *x g* for 1 minute. This step was repeated for a total of 60μL of elution volume. RNA concentration was determined by nanodrop and DNase 1 treated (NEB, Cat. # M0303L). DNase 1 (4 units) was added to each sample and let to incubate at 37°C for 30 minutes. RNA was re-purified by adding 350 μL of RLT buffer containing BME, followed by 250 μL of 100% ethanol and mixed by pipetting. The solution was transferred to an RNeasy Mini Spin column and washed and eluted as above. The concentration of each sample was determined by nanodrop. RNA was confirmed free of DNA contamination by running a qPCR for the *BT1155* gene of *B. theta* (see Supplemental Table 1) using an in-house qPCR mix (see below) on the purified RNA prior to cDNA synthesis, and were considered DNA-free when a Ct value greater than our background value of 30 was obtained (72, 73).

Reverse transcription was performed using an adapted SuperScript III First-Strand Synthesis System (Invitrogen, Cat. # 18080-051) to generate cDNA. In brief, 1 μg of RNA was incubated with 10 mM dNTP mix (1 μL/sample) and random hexamers (200 ng/sample) in a final volume of 13 μL at 65°C for 5 minutes followed by 5 minutes on ice. Each sample was spun down and first strand buffer (1X, 4 μL/sample), 0.1M DTT (1 μL/sample), SuperScript III enzyme (200 units/sample), and water (1 μL/sample) were added to a final volume of 20 μL. The sample was mixed by pipetting gently and the tube was spun down prior to incubating for 1 hour at 60°C. The reaction was halted by inactivating the enzyme at 70°C for 15 minutes. A 1-to-1 conversion of RNA to cDNA was assumed and each sample was diluted with water to 5 ng/μL and stored at -20°C until use.

The abundance of each target transcript (*BT1154-BT1161*) was quantified using an in-house qPCR mix as described in (72, 73) (see Supplemental Table 1 for qPCR primer sequences). In brief, each 20 μL reaction contained 1X SYBR Green 1 (Lonza, Cat. # 50513), 1X Thermopol Reaction Buffer (NEB, Cat. # B9004S), 125 μM dNTPs, 2.5 mM MgSO4, 500 nM of gene specific primers, and 0.5 units Hot Start *Taq* Polymerase (NEB, Cat. # M0495L), and 10 ng of template cDNA. 16S rRNA was used as a housekeeping gene for each sample, and a water control was run for each primer condition. A Ct value greater than our background value of 30 was considered non-detectable (72, 73). Each sample was run in duplicate and the Ct values were averaged. The ΔCt values were calculated by normalizing the gene of interest Ct value to the 16S rRNA housekeeping gene value for each sample. Next, the Δ/ΔCt value was calculated by normalizing the ΔCt value for *BT1160* deletion samples to the ΔCt value of the parent strain (Δ*tdk*). Fold change was calculating using the 2^Δ/ΔCt^ method.

### *B. theta* mutant IL-10 induction following NAD+ supplementation

NAD+ (Millipore Sigma, Cat. # 481911) was reconstituted in Milli-Q water to make a stock solution of 50 mg/mL (75.37 mM) and sterile filtered using a 0.22 μm PVDF filter (Millex, Cat. # SLGVR33RS). The stock solution was aliquoted and stored at -20°C until use. NAD+ was either added directly to TYG bacterial growth media at the start of the bacterial culture, or added directly to the RPMI media in which the splenocytes were cultured. NAD+ was added to either TYG or RPMI at a final concentration of 0 μM, 1 μM, 10 μM, or 100 μM. For NAD+ supplementation into the bacterial culture media, NAD+ was added to TYG and then inoculated with either *B. theta*^Δ*tdk*^, *B. theta*^Δ*tdk*Δ*BT1160_*trunc^, or *B. theta*^Δ*tdk*Δ*BT1155-BT1160*^, or TYG control media. Cultures were grown up anaerobically overnight, normalized by OD_600_, and the conditioned media was prepared as described above. The conditioned media for each *B. theta* mutant at each dose of NAD+ (0 μM, 1 μM, 10 μM, 100 μM), was tested for its ability to induce IL-10 by wild-type splenocytes (as described above) at three different doses (1%, 3%, or 5% v/v). For NAD+ supplementation directly onto wild-type splenocytes, NAD+ was added to RPMI at a final concentration of 0 μM, 1 μM, 10 μM, 100 μM. Wild-type splenocytes were plated in NAD+ supplemented RPMI followed by stimulation with conditioned media from either *B. theta*^Δ*tdk*^, *B. theta*^Δ*tdk*Δ*BT1160_*trunc^, or *B. theta*^Δ*tdk*Δ*BT1155-BT1160*^, or TYG control media to which no NAD+ had been added at three different doses (1%, 3%, 5%, v/v) and IL-10 induction was measured by ELISA as described above.

### OMV comparative proteomics using liquid chromatography with tandem mass spectrometry (LC-MS/MS)

Outer membrane vesicles were isolated from *B. theta*^Δ*tdk*Δ*BT1160_*trunc^, *B. theta*^Δ*tdk*Δ*cps*1-8Δ*BT1160_*trunc^, and their corresponding parent strains (*B. theta*^Δ*tdk*^ and *B. theta*^Δ*tdk*Δ*cps*1-8^) grown to stationary phase in TYG. Conditioned media from three 250 mL cultures of each mutant was made as described above. Each 250 mL batch of conditioned media was divided between six 39 mL QuickSeal Ultracentrifugation tubes (Beckman Coulter, Cat. # 342414) and OMVs isolated by ultracentrifugation at 100,000 *x g* (as described above). The remaining conditioned media was kept to test its ability to induce IL-10 using the *in vitro* splenocyte assay (described above) to confirm that the conditioned media used in these measurements displayed the expected immune-stimulatory capacity. OMV pellets were resuspended and combined in a total of 1 mL sterile PBS to wash, and put through another round of ultracentrifugation. Washed OMV pellets were resuspended in 300 μL sterile Milli-Q water and frozen at -80°C. An aliquot was taken to test their ability to induce IL-10 to confirm that the OMV used in these measurements displayed the expected immune-stimulatory capacity, and were further assessed by ZetaView (as describe below) to ensure OMVs were successfully isolated.

Prior to LC-MS/MS, samples were lyophilized and then lysed in 200 μL Radio-Immunoprecipitation Assay (RIPA) buffer using a probe sonicater, and protein concentration was measured by BCA assay. A 100 μg aliquot of each sample was precipitated with acetone overnight, and the pellets dried and reconstituted in 6M urea/Tris buffer. Samples were then reduced with 1 μL of 200 mM dithiothreitol, alkylated by 4 μL of 200 mM iodoacetamide, then digested with 4 μL of a trypsin/Lys-C Mix (0.5 μg/uL) (Promega # V5071) using a two-step in-solution digestion overnight at 37°C. Digestion was terminated by the addition of trifluoracetic acid to a final concentration of 0.5-1% v/v. The samples were then desalted using Sep-Pak C18 cc Vac Cartridge (WAT054955) and reconstituted in 30 μL 0.1% v/v formic acid.

Prepared samples were run on a LC-MS Dionex Ultimate 3000 nano-flow high performance liquid chromatography (HPLC) interfacing with a ThermoScientific Fusion Lumos mass spectrometer system. The HPLC system used an Acclaim PepMap 100 precolum (75 μm x 2 cm, C18, 3 μm, 100 A) followed by an Acclaim PepMap RSLC analytical column (75 μm x 15 cm, C18, 2 μm, 100 A). 5 μL volumes of the extract were injected and the peptides eluted from the column by an acetonitrile/0.1% v/v formic acid gradient at a flow rate of 0.3 μL/min were introduced into the source of the mass spectrometer online. The micro-electrospray ion source is operated at 2.5 kV. The digest was analyzed using the data dependent multitask capability of the instrument acquiring full scan mass spectra to determine peptide molecular weights and product ion spectra to determine amino acid sequence in successive instrument scans. The data were searched against the *B. theta* (VPI-5482) UniProtKB protein database with the program Andomeda (4782 sequences). LC-MS spectra intensities for identified proteins were normalized and expressed as label free quantity (LFQ) intensities calculated in the program MaxQuant version 2.0.1.0. The data matrix was uploaded into Perseus and proteins that were not found in at least 2 samples were discarded from the analyses. Zero intensity values were imputed by replacing intensities with a normal distribution. The Δ*tdk*/Δ*tdk*Δ*BT1160_*trunc LFQ ratio (fold change) was calculated on averaged LFQ intensities and the p-value calculated using Student’s T-test.

### OMV quantification and normalization

The concentration of OMVs isolated from *B. theta* mutants lacking *BT1160, BT1155-BT1159,* and *BT1155-BT1160* and their parent strain were determined using Qubit (measures protein concentration) or ZetaView (a nanoparticle analyzer with Particle Metrix software) (74). OMVs were isolated from 1 mL aliquots of conditioned media from *B. theta*^Δ*tdk*^ and mutant strains as described above. The concentration of isolated OMVs were measured by Qubit and Zetaview. For Qubit (Invitrogen, Cat. # Q33211), the OMVs from 1 mL of conditioned media were concentrated 50X by resuspending in 20 μL of sterile filtered PBS. The concentration was measured in μg/mL and then the 1X original concentration of OMVs was calculated. For ZetaView, OMVs isolated from 1 mL of conditioned media were resuspended in a volume of sterile filtered PBS equivalent to the original volume from which they were obtained (as described above) and 1 mL was injected into the ZetaView. Once the concentration was determined for both the mutants and parent strain, OMVs were isolated from the same batch of conditioned media and resuspended in PBS at either the original volume (for *B. theta*^Δ*tdk*^) or, for mutants whose OMV concentration was lower (example: *B. theta*^Δ*tdk*Δ*BT1160*^), in a volume that resulted in the same concentration of OMVs as *B. theta*^Δ*tdk*^. Concentrated OMVs from the mutants and OMVs from the parent strain were then tested for their ability to induce IL-10 using the *in vitro* IL-10 assay described above. The same percentage of isolated OMVs, from both the mutant and parent strain, were added to the splenocytes.

### Transmission electron microscopy (TEM) of OMVs

50 mL cultures of *B. theta*^Δ*tdk*^ and *B. theta*^Δ*tdk*Δ*BT1160_*trunc^ were grown in TYG anaerobically overnight to stationary phase at 37°C and the OD_600_ was measured. Mutants were normalized to a final OD of 2.95 using TYG and the conditioned media was isolated as described above. Briefly, cultures were spun at 7000 *x g* for 5 minutes and the supernatant was filtered through a 0.2 μm PES filter (MilliQ Steriflip, Cat. #SCGP00525). 1 mL aliquots of conditioned media were removed and frozen at -20°C to test the mutant and parent strains ability to induce IL-10 using the assay described above to confirm that those OMV that were imaged showed the expected immune-stimulatory characteristics. The conditioned media from each mutant was then concentrated from 50 mL to 1 mL using 100 kDa size cut off filters (Vivaspin20, Cat. # 28932363). OMVs remain in the larger than 100 kDa top fraction and were then subject to OMV isolation using the ultracentrifugation steps described above. The washed pellet containing the OMVs was resuspended in 100 μL of PBS, the solution was turbid, and frozen at -20°C until use.

OMVs were prepared for TEM by negative staining. 5 μL of each OMV sample was added to the carbon-coated copper 400 mesh grid and incubated at room temperature for 10 minutes. The samples were washed with 5 drops of 5 mM Tris buffer (pH 7.1) and then with 5 drops of distilled water. The specimen was stained with 5 drops of 2% w/v uranyl acetate. The grid was allowed to dry completely for several hours or overnight. The prepared specimens were visualized with an electron microscope operated at 80 kV.

### Statistical analysis

Statistical analyses were performed in GraphPad Prism version 9.0.0. Details of the specific statistical tests used is included in the appropriate figure legends.

## Results

### *B. thetaiotaomicron* secretes immunomodulatory factors that are sensed via a TLR2-MyD88 axis

*B. thetaiotaomicron* (*B. theta*) has been established as a potent immunomodulatory microbiome member (9, 34, 35, 45, 53). We sought to define the pathways that underlie the capacity of *B. theta* to mediate such effects. Several bacterial members of the microbiome have been shown to stimulate the production of IL-10 via secreted factors (14, 15, 31, 33). To determine if *B. theta* also stimulates immune responses via secreted factors, we treated unfractionated splenocytes from wild-type mice with either heat-inactivated *B. theta* (strain VPI-5482) cell pellets (OD_600_=1.0 per 5X10^5^ splenocytes), conditioned media (sterile-filtered TYG media conditioned by growth of *B. theta*; 1% v/v), or sterile TYG bacterial culture media for two days, and assessed the production of the cytokine IL-10 in culture supernatants by ELISA. Both bacterial pellets and conditioned media stimulated significant production of IL-10, demonstrating that *B. theta* drives production of IL-10 and that this can be mediated by factors that are secreted/released from the cell (Figure 1A). Furthermore, these findings were replicated using murine bone-marrow derived macrophages (BMDMs) (75) suggesting that this response is not limited to splenocytes (Supplemental Figure 1A). As with many microbes/microbial products, both cell pellet and conditioned media also induced significant production of IL-6 (Supplemental Figure 1B), often associated with inflammation (76), but notably, IL-6 is also required for the development of bacterial-reactive anti-inflammatory Tregs in the intestine (77, 78). Thus, IL-6 also forms part of a loop that sustains mutualism. As *B. theta* elicits Treg development and does not elicit inflammation in wild-type mice (34, 45), induction of IL-10 and IL-6 likely both contribute to its anti-inflammatory properties, and thus we focused our remaining analyses on its ability to induce IL-10. Outer membrane vesicles (OMVs) are spherical vesicles that are released by many Gram-negative bacteria. *B. theta* produces OMVs that have been posited to represent the dominant process through which it interacts with the host (45). Moreover, OMVs from the related bacterium, *B. fragilis*, have been shown to induce IL-10 (33). To determine if OMVs secreted into the conditioned media by *B. theta* can stimulate IL-10, we isolated OMVs from stationary phase cultures via ultracentrifugation, resuspended them in a volume of PBS equivalent to that from which they were prepared, and used them to stimulate splenocytes for two days (1% v/v). These OMVs also induced significant production of IL-10 relative to TYG culture media control which had been passed through the same process (Figure 1B), although notably, the level of induction was slightly less than that seen with unfractionated conditioned media. Importantly, while all isolated batches of OMVs were capable of inducing IL-10, we did observe variability in the proportion of IL-10 induced compared to unfractionated conditioned media among different batches. This suggests that factors that are secreted into the conditioned media rather than packaged into OMV could also mediate immunomodulatory function or that some facet of the isolation process leads to destruction of the OMV and the release of these factors into the conditioned media.

**Figure 1.**
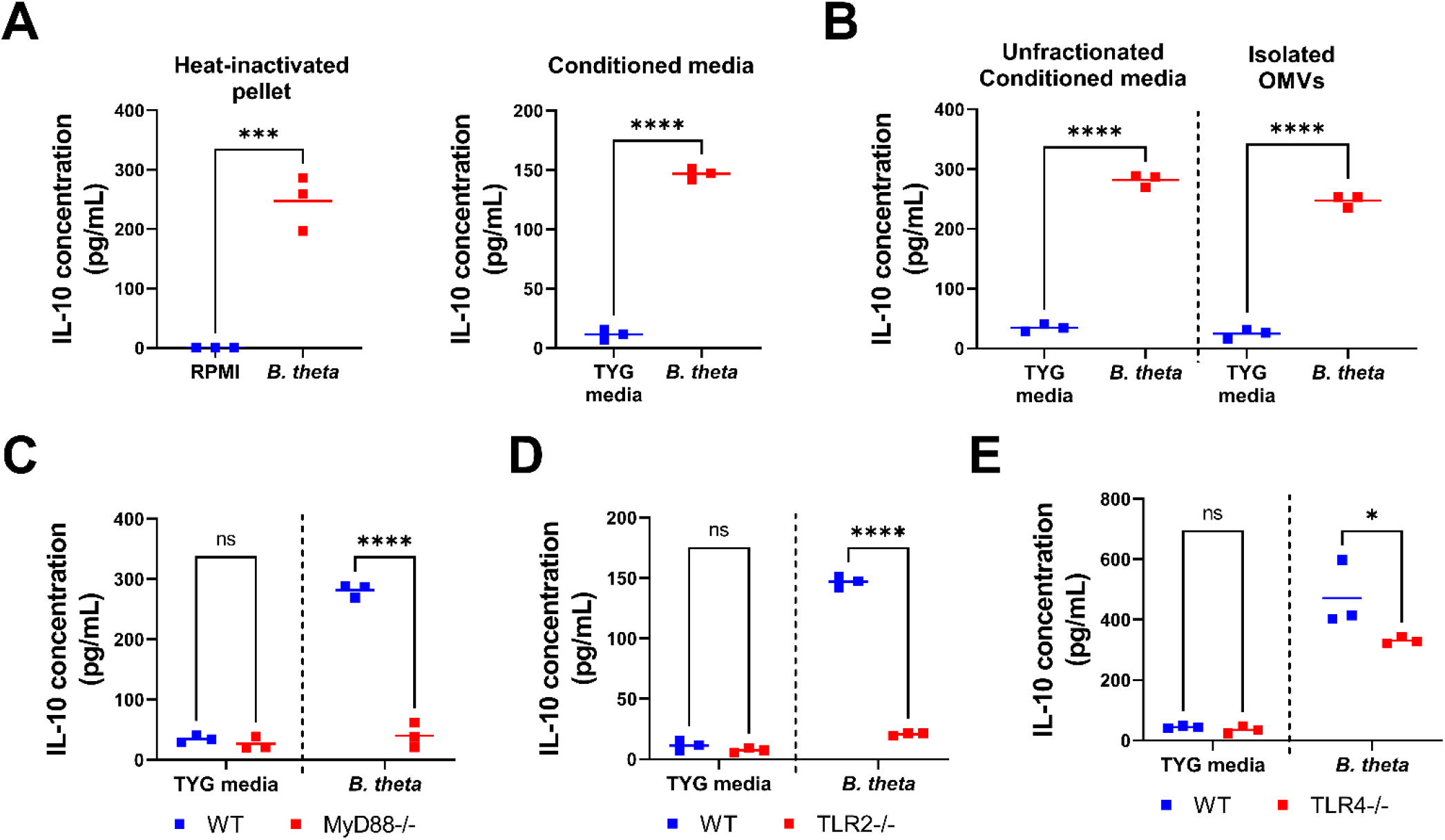
*B. theta* secretes immunomodulatory factors that are sensed via a TLR2-MyD88 axis. **(A)** Unfractionated wild-type splenocytes were stimulated with *B. theta*^VPI-5482^ heat-inactivated pellet (OD =1.0) or *B. theta*^VPI-5482^ conditioned media (1% v/v), or appropriate media control for 2 days and IL-10 was assessed in the supernatant by ELISA. **(B)** Wild-type splenocytes were stimulated with *B. theta*^VPI-5482^ unfractionated conditioned media or TYG media (1% v/v), or isolated OMVs from the conditioned media or TYG control media that underwent the same process as that for OMV isolation (resuspended in an equivalent volume of PBS to that of the conditioned media from which they were isolated) (1% v/v). Samples were incubated for 2 days and IL-10 was assessed in the supernatant by ELISA. **(C-E)** Wild-type and MyD88^-/-^ **(C)** TLR2^-/-^ **(D)** or TLR4^-/-^ **(E)** splenocytes were stimulated with *B. theta*^VPI-5482^ conditioned media or TYG media control (1% v/v) for 2 days and IL-10 was assessed in the supernatant by ELISA. Data points represent an independent technical replicate and horizontal lines show the mean **(A-E)**. Graphs are representative of 4 experiments **(A-E)**. Statistical significance was determined using Student’s T test: **(A, B)**, and two-way ANOVA with Sidak’s post hoc test, comparisons to wild-type **(C-E)**. ns (not significant) P≥0.05; **\*\***P≤0.01, ***P≤0.001, ****P≤0.0001.

*B. fragilis* produces a capsular polysaccharide (CPS), polysaccharide A (PSA), that can be packaged into OMVs and drives IL-10 induction in a TLR2-dependent manner (33). To determine if there were parallels with how *B. theta* induced IL-10, such as a dependency on TLRs, we first assessed the role of the signaling adaptor MyD88, that mediates most TLR signaling pathways (79), in responsiveness to *B. theta*. To do so, we stimulated wild-type or MyD88^-/-^ splenocytes with conditioned media and assessed IL-10 in culture supernatant after two days, revealing that *B. theta* conditioned media promotes IL-10 induction via MyD88 (Figure 1C). These data suggest that *B. theta* produces a factor recognized by a TLR. We next investigated the role of TLR2 in *B. theta* elicited induction of IL-10 by stimulating wild-type and TLR2^-/-^ splenocytes with *B. theta* conditioned media. As with *B. fragilis*, TLR2 was required for responsiveness to *B. theta* as TLR2^-/-^ splenocytes produced significantly less IL-10 than wild-type splenocytes in response to *B. theta* conditioned media (Figure 1D). Stimulation of BMDMs generated from the bone-marrow of wild-type or TLR2^-/-^ mice, also showed a role for TLR2 in producing IL-10 in response to *B. theta* (75) (Supplemental Figure 1C), confirming that the requirement for TLR2 was not limited to the splenocyte assay. Conditioned media from an additional isolate of *B. theta* (strain 0940-1) also induced IL-10 in a predominantly TLR2-MyD88-dependent manner (Supplemental Figure 1D, E), suggesting that TLR2 is a sensor for *B. theta* as a species, in keeping with a recent report showing that a large panel of distinct *Bacteroides* are sensed by this PRR (80). While the majority of the IL-10 induced by *B. theta* conditioned media was dependent on TLR2, we aimed to investigate if the residual *B. theta* driven IL-10 in TLR2^-/-^ splenocytes was driven through another common PRR, such as TLR4, a sensor of LPS (81, 82). Stimulation of wild-type and TLR4^-/-^ splenocytes revealed a moderate dependency of *B. theta* induced IL-10 on TLR4 (Figure 1E), however, this was only observed in ∼50% of batches tested (data not shown). Moreover, conditioned media from the *B. theta* strain 0940-1 also showed a moderate decrease in IL-10 induction in TLR4^-/-^ splenocytes (Supplemental Figure 1F). Collectively, these data suggest that while TLR4 may play a role in stimulating IL-10 production in response to *B. theta,* a TLR2-MyD88 axis is the predominant pathway through which *B. theta* derived immunomodulatory factors are sensed. TLR2 can coordinate activation of the immune system in concert with the PRR Dectin-1, and the dual engagement of TLR2 and Dectin-1 by *B. fragilis* derived PSA drives IL-10 production (30, 83). However, distinct from *B. fragilis*, we found that the ability of secreted factors produced by *B. theta* to induce IL-10 is not dependent on Dectin-1 (Supplemental Figure 1G). These data suggest the existence of a novel factor in *B. theta* that induces IL-10.

### Gene *BT1160* is required for *B. thetaiotaomicron* mediated induction of IL-10

To further investigate how *B. theta* induces the production of IL-10 we aimed to identify genes that were required for its IL-10 inducing capacity. To do so, we used an arrayed transposon mutagenesis library of 2,009 *B. theta* transposon mutants, which had previously been generated on the acapsular genetic background (*B. theta^ΔtdkΔcps^*^1-8^) (35), allowing the assessment of individual mutations on IL-10 inductive capacity. Splenocytes were stimulated with the conditioned media from individually-grown *B. theta^ΔtdkΔcps^*^1-8^ transposon mutants (1% v/v per well) for two days (note, to facilitate testing of a large batch of mutants, the conditioned media was centrifuged to remove cellular debris but was not sterile filtered). We then assessed IL-10 induction in culture supernatants by ELISA, and normalized the values for each transposon mutant to the positive control, Pam3CSK4. Several mutants displayed impaired IL-10 induction as evidenced by a lower IL-10/Pam3CSK4 ratio (Figure 2A). We chose one mutant for follow-up (mutant from well H4 of plate 19; 19_H4), and validated its impaired ability to induce IL-10 by generating an independent batch of sterile-filtered conditioned media which confirmed its impaired ability to induce IL-10 (Figure 2B). Next we mapped the location of the transposon insertion, and found that transposon mutant 19_H4 had a disrupted, out-of-frame insertion in gene *BT1160*, which is predicted based on gene homology to encode subunit A (*nqrA*) of the Na+-transporting NADH:ubiquinone oxidoreductase (NQR complex). The NQR complex is comprised of six subunits encoded by a six gene cluster (*nqr* locus, genes *BT1155-BT1160*) (Figure 2C). The NQR complex has been characterized in other bacteria, including *B. fragilis*, and is found within the inner membrane, responsible for converting NADH to NAD^+^ and shuttling electrons down the electron transport chain, to generate ATP, while pumping sodium from the cytosol into the periplasm to create an ion gradient and converting fumarate to succinate (55–57). To validate the role of this gene we sought to create engineered in-frame deletions of gene *BT1160*. First, we generated a truncated knock-out of gene *BT1160* (Δ*BT1160_*trunc) on both the acapsular (Δ*tdk*Δ*cps*1-8) and wild-type (Δ*tdk*) backgrounds. We engineered this truncated deletion of gene *BT1160* with the goal of limiting modifications to the downstream genes of the *nqr* locus as the predicted promotor for the *nqr* locus is directly upstream of gene *BT1160*. The truncated deletion left the first 8 nucleotide bases of gene *BT1160*— including the start codon of *BT1160*—intact. However, the same reading frame encodes a stop codon prior to gene *BT1159*, therefore, downstream genes are not shifted out of frame. This deletion of gene *BT1160* resulted in significant impairment in IL-10 induction by *B. theta* compared to the parent strain, on both the wild-type and acapsular backgrounds (Figure 2D). Therefore, the impact of gene *BT1160* on the ability of *B. theta* to induce IL-10 was not limited to the acapsular background. It should be noted that across multiple independent assessments (>30) of IL-10 induction using this mutant, we observed rare instances where no significant differences were detected between mutant and wild-type. The ability of *B. theta* conditioned media to induce IL-10 from BMDMs was also shown to be dependent on gene *BT1160* (Supplemental Figure 1H). Thus, our screen identified gene *BT1160* as a *bona fide* regulator of IL-10 induction in *B. theta*. Although we observed statistically significant impacts on growth following deletion of gene *BT1160*, the magnitude of these effects was minimal, and the mutant reached equivalent levels of growth as wild-type in stationary phase (Supplemental Figure 1I) (note, the Δ*BT1160_*trunc mutant grew significantly more slowly than parent strain directly from frozen stocks as has been reported for mutants in this pathway in *B. fragilis* (*56*), however, parent strain-levels of growth were observed when sub-cultured, and thus in all assays, the cultures used were subcultures from initial overnight cultures that were grown to stationary phase).

**Figure 2.**
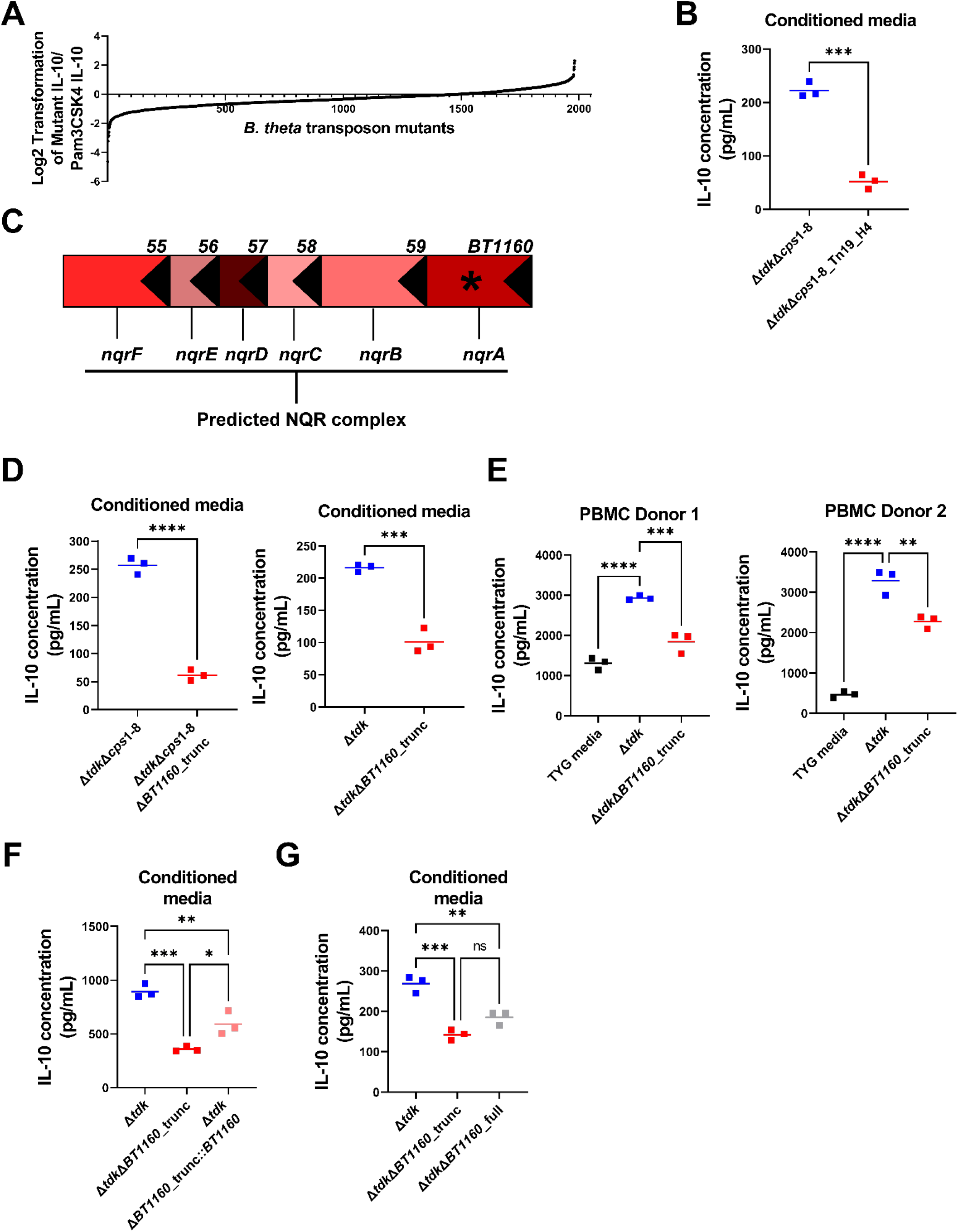
Gene BT1160 is required for B. theta mediated induction of IL-10. **(A)** Wild-type splenocytes were stimulated with conditioned media from a library of 2,009 individual *B. theta*^Δ^*^tdk^*^Δ^*^cps^*^1-8^ transposon mutants (1% v/v) or Pam3CSK4 for 2 days and IL-10 levels were measured by ELISA. For each plate, IL-10 levels were normalized to the IL-10 level induced by the positive control Pam3CSK4. Data shows the Log2 transformed value normalized to the positive control Pam3CSK4 of the 1982 mutants whose growth was higher than background control media. **(B)** Wild-type splenocytes were stimulated with conditioned media from *B. theta*^Δ^*^tdk^*^Δ^*^cps^*^1-8^ or transposon mutant 19_H4 (1% v/v) for 2 days and IL-10 was assessed by ELISA. **(C)** The transposon insertion in transposon mutant 19_H4 was mapped to gene *BT1160* that is predicted to encode the first subunit of the NQR complex. **(D)** Wild-type splenocytes were stimulated with conditioned media (1% v/v) from *B. theta* harboring a truncated deletion of gene *BT1160* (acapsular background, left; wild-type background, right), or appropriate parent strain, for 2 days, and IL-10 levels were measured by ELISA. **(E)** Human PBMCs from two healthy donors were stimulated with conditioned media (3% v/v) from *B. theta* harboring a truncated deletion of gene *BT1160* in the wild-type background, parent strain (Δ*tdk*), or TYG media for 2 days, and IL-10 levels were measured by ELISA. **(F)** Wild-type splenocytes were stimulated with conditioned media (1% v/v) from *B. theta*^Δ^*^tdk^*, *B. theta*^Δ^*^tdk^*^Δ^*^BT1160_^*trunc, or *B. theta*Δ*^tdk^*^Δ^*^BT1160_^* trunc complemented with *BT1160* (*B. theta*Δ*^tdk^*^Δ^*^BT1160_^*^trunc*::BT1160*) for 2^ days, and IL-10 levels were measured by ELISA. **(G)** Wild-type splenocytes were stimulated with conditioned media (1% v/v) from *B. theta* harboring a truncated deletion of gene *BT1160* (red), a full deletion of gene *BT1160* (grey) (Δ*tdk* background), or the parent strain *B. theta*^Δ^*^tdk^* (blue) for 2 days, and IL-10 levels were measured by ELISA. Individual data points show mean values **(A)** or individual technical replicates **(B, D-G)**, and horizontal lines represent the mean. Graphs are representative of at least 4 individual experiments **(B, D, F, and G)** and 1 individual experiment **(A, E)**. Statistical significance was determined by Student’s T-test: **(B, D)**, and one-way ANOVA with Dunnett’s post-hoc test, comparisons to *B. theta*^Δ*tdk*^ or between all **(E-G)**. ns (not significant) P≥0.05, **\***P<0.05, **P≤0.01, ***P≤0.001, ****P≤0.0001.

*B. theta* has been reported to stimulate IL-10 production from human peripheral blood mononuclear cells (PBMCs) (53). To determine if *BT1160* played a role in the immunomodulatory function of *B. theta* on human cells, we stimulated PBMCs from two different healthy donors with the conditioned media from wild-type (Δ*tdk*) *B. theta* or the truncated *BT1160* deletion mutant and found that as with murine cells, the induction of IL-10 in human PBMCs also required *BT1160* (Figure 2E). Complementation of *BT1160 in trans* by introduction of the full-length *BT1160* gene and the promotor region into the *att2* (tRNA^ser^) site, partially restored the ability of *B. theta* to induce IL-10, compared to wild-type (Figure 2F), with notable variance between experiments (Supplemental Figure 2A). Next, we generated a complete in-frame deletion of gene *BT1160* on the Δ*tdk* background, that also significantly diminished the ability of *B. theta* to induce IL-10 compared to the parent strain, but did not impair IL-10 induction as greatly as the truncated genetic mutant (Figure 2G). Like the complementation of gene *BT1160* in the truncated mutant, the full deletion of *BT1160* displayed variation in IL-10 induction between experiments (data not shown). These data show that disruption of gene *BT1160* significantly impairs *B. theta* mediated IL-10 induction. However, given the differences between the two deletion mutants, the variance in the magnitude of effects between independent experiments, and the partial impact of complementation, the effects observed in the *BT1160* deletion mutants may be attributable to other genes that are impacted in these mutants, likely the other genes in the *nqr* locus.

### The NQR complex coordinates IL-10 induction by *B. thetaiotaomicron*

The NQR complex is encoded by a bacterial locus (*nqr*) made up of six genes, genes *BT1155*-*BT1160* (Figure 3A). Genes *BT1155-BT1159* code for subunits B-F (*nqrB-F*) in the NQR complex and are directly downstream of *BT1160* (Figure 3A). To determine if deletion of gene *BT1160* impacted expression of these downstream genes, we performed RT-qPCR to assess gene expression at mid-log phase in both the truncated and full genetic deletion mutants. These data showed that both *BT1160* mutants displayed a reduction in expression of downstream genes *BT1156-BT1159* within the *nqr* locus compared to wild-type *B. theta* (Δ*tdk*) (Figure 3B), suggesting that the reduced IL-10 induction by these mutants may be attributable, in part, to altered expression of these genes (as expected, expression of gene *BT1160* was not detectable in either mutant; not shown). To determine if alterations in expression of other *nqr* genes could also impact IL-10 induction, we engineered *B. theta* mutants lacking either the entire *nqr* locus (Δ*BT1155-BT1160*) or the entire locus except *BT1160* (Δ*BT1155-BT1159*). Both mutants were significantly impaired in their capacity to induce IL-10 from splenocytes across a range of doses (0.5%, 1%, and 3% v/v) (Figure 3C, D and Supplemental Figure 2B; note, the IL-10 values for *B. theta*^Δ*tdk*^ in Figure 3C and D represent the same data), and did so across all experiments, reinforcing the notion that the complex as a whole is required for IL-10 induction. Furthermore, the induction of IL-10 by *B. theta* conditioned media, and its dependence on *nqr* genes, was observed in cells from both male and female mice, suggesting this pathway is not limited to one biological sex (Supplemental Figure 3A). Importantly, the expression of genes *BT1154* and *BT1161*, which lie directly downstream and upstream of the *nqr* locus, were not significantly altered by the deletion of *BT1160* and a control deletion mutant of gene *BT1161* has no discernible impact on IL-10 induction (Figure 3B, E). These data suggest that (i) disruption of the *nqr* locus impairs the ability of *B. theta* to induce IL-10, but that deletion of a gene directly upstream of the locus does not, and that (ii) *BT1160* in the absence of other *nqr* genes is not sufficient for induction of IL-10, and thus the impact of deleting gene *BT1160* may be mediated by alterations on other genes within the *nqr* locus.

**Figure 3.**
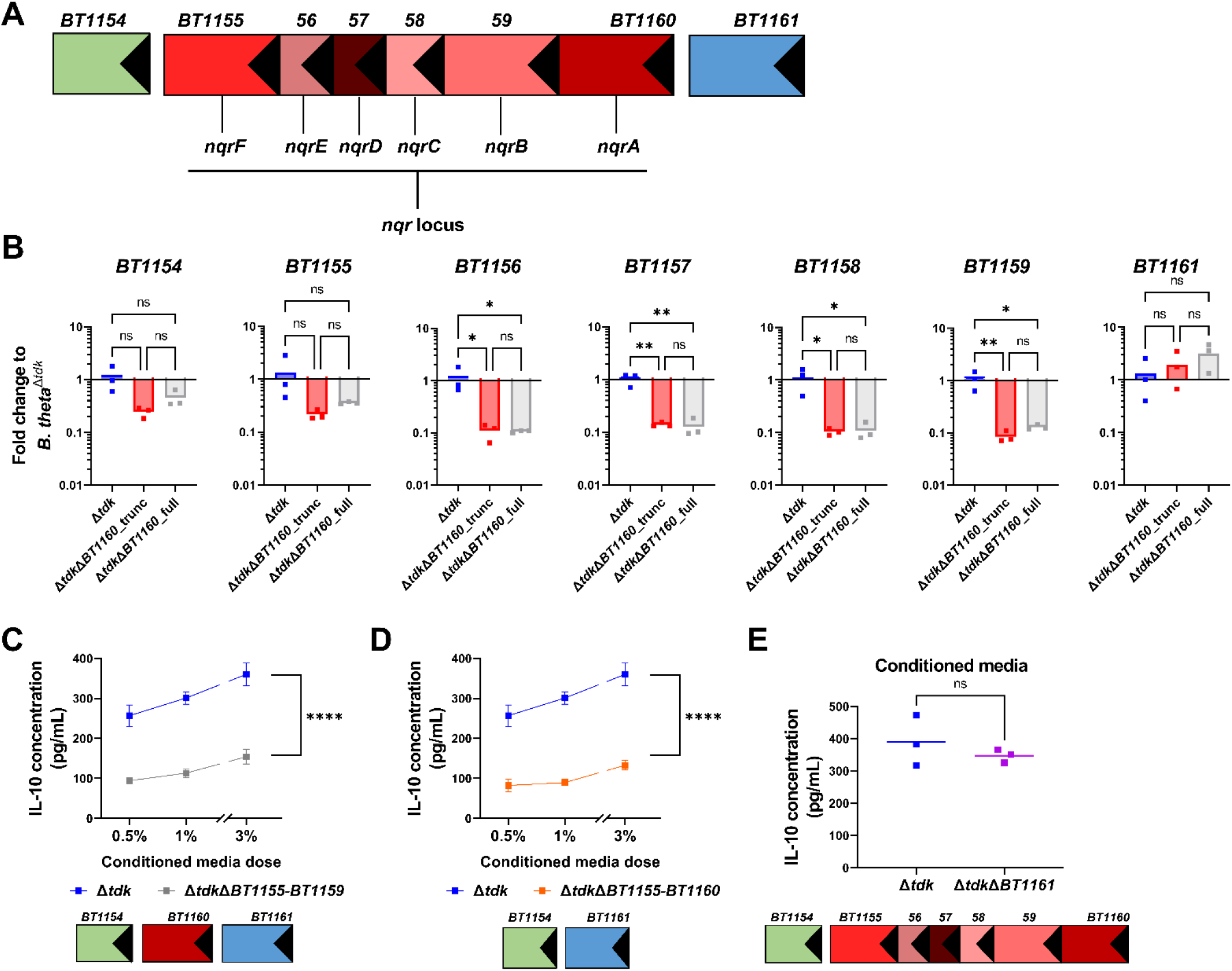
*B. theta* NQR complex promotes IL-10 induction. **(A)** Schematic of the gene locus that encodes the NQR complex and flanking genes. **(B)** The expression (2^Δ/Δ^ ^Ct^ ^values^) of *nqr* locus genes *BT1155-BT1159* and control flanking genes (*BT1154* and *BT1161*). *B. theta*^Δ*tdk*^, *B. theta* ^Δ^*^tdk^*^Δ^*^BT1160_^*^trunc^, and *B. theta* ^Δ^*^tdk^*^Δ^*^BT1160^*^_full^ were grown *in vitro* in TYG media and cells were harvested at mid-log growth, and RNA expression profiles were assessed by RT-qPCR with the *B. theta* 16S rRNA gene as the housekeeping gene. **(C-E)** Wild-type splenocytes were stimulated with different doses of conditioned media (0.5%, 1%, 3% v/v) from *B. theta* harboring deletions of genes *BT1155-BT1159* **(C)**, or genes *BT1155-BT1160* **(D)**, and conditioned media (1% v/v) from *B. theta* harboring a deletion in gene *BT1161* **(E)**, on the *B. theta*^Δ*tdk*^ background for 2 days, and IL-10 levels were measured by ELISA. Data points represent the mean of technical replicates of independent biological replicates **(B)** or independent technical replicates **(E)**, or the mean of three technical replicates **(C, D)** and horizontal bars represent the mean **(B, E)**, and error bars shown the standard deviation **(C, D)**. Graphs are representative of a single experiment **(B)** or 4 experiments **(C-E)**. Note, the IL-10 values for *B. theta*^Δ*tdk*^ in C and D represent the same data. Statistical significance was determined using one-way ANOVA with Dunnett’s post hoc test and comparisons between all groups: **(B)**, and Student’s T test **(E)** on area under the curve: **(C-D)**. ns (not significant) P≥0.05, **\***P<0.05, **P≤0.01, ***P≤0.001, ****P≤0.0001.

### NQR complex NAD+ regeneration is not essential for IL-10 induction

We next aimed to uncover how the NQR complex could impact induction of IL-10. The NQR complex resides within the bacterial inner membrane and plays a role in maintaining the energetics of the cell (55–57). The complex regenerates NAD+ (an electron acceptor) from NADH and shuttles an electron to downstream members of the electron transport chain. Disruption of the NQR complex whether it be through deletion of the gene predicted to encode subunit A of the six subunit complex (Δ*BT1160*_trunc) or the entire *nqr* locus (Δ*BT1155-BT1160*) could thus alter these downstream processes. Importantly, extracellular NAD+ has recently been reported to directly modulate immune function whether it be through direct binding to purinergic receptors or acting as a co-factor for enzymes that can bind TLRs (84–86). Therefore, alterations in the regeneration of NAD+ could account for the inability of *B. theta* conditioned media to induce IL-10 when the NQR complex is disrupted or deleted. We therefore aimed to investigate whether (i) NAD+ directly impacts the ability of *B. theta* to produce immunomodulatory factors, or (ii) if NAD+ released by *B. theta* into conditioned media play a role in stimulating the induction of IL-10. To test this we supplemented both bacterial cultures and splenocyte culture media with NAD+ during growth and stimulation with conditioned media respectively. Firstly, we addressed whether a lack of regenerated NAD+ in *nqr* mutants altered the immunomodulatory potential of *B. theta* conditioned media compared to the parent strain. Based on published literature exogenous NAD+ was supplemented into TYG bacterial medium to a final concentration of 0 μM, 1 μM, 10 μM, or 100 μM (87). The *B. theta* cultures supplemented with or without NAD+ were grown to stationary phase and conditioned media was generated and tested for its ability to induce IL-10. Supplementation of NAD+ did not restore the ability of *B. theta*^Δ*tdk*Δ*BT1160_*trunc^ or *B. theta*^Δ*tdk*Δ*BT1155-BT1160*^ conditioned media to induce IL-10, with the *nqr* mutant’s conditioned media remaining unable to induce IL-10 compared to *B. theta*^Δ*tdk*^ regardless of the dose of NAD+ used (Figure 4A). Additionally, we addressed whether a potential lack of NAD+ in the *B. theta* conditioned media upon disruption or deletion of the NQR complex directly influenced the ability of the mutant’s conditioned media to induce IL-10, compared to conditioned media from the parent control. *B. theta* conditioned media from the different NQR complex mutants or parental control was added to wild-type splenocytes with or without NAD+ supplementation to a final concentration of 0 μM, 1 μM, 10 μM, or 100 μM. Addition of NAD+ directly to the splenocyte culture also did not restore the ability of *B. theta*^Δ*tdk*Δ*BT1160_*trunc^ or *B. theta*^Δ*tdk*Δ*BT1155-BT1160*^ conditioned media to induce IL-10 compared to *B. theta*^Δ*tdk*^ (Figure 4B). These data suggest that provision of exogenous NAD+ is not sufficient to restore IL-10 induction, and that impaired NAD+ regeneration by *B. theta* mutants with a defective NQR complex is not directly responsible for their reduced ability to stimulate IL-10 production, however, we cannot completely rule out a role for NAD+ that was not readily revealed in our system. Additionally, these data suggest that the NQR complex may play an indirect role, influencing further downstream processes that alter the ability of *B. theta* to induce IL-10.

**Figure 4.**
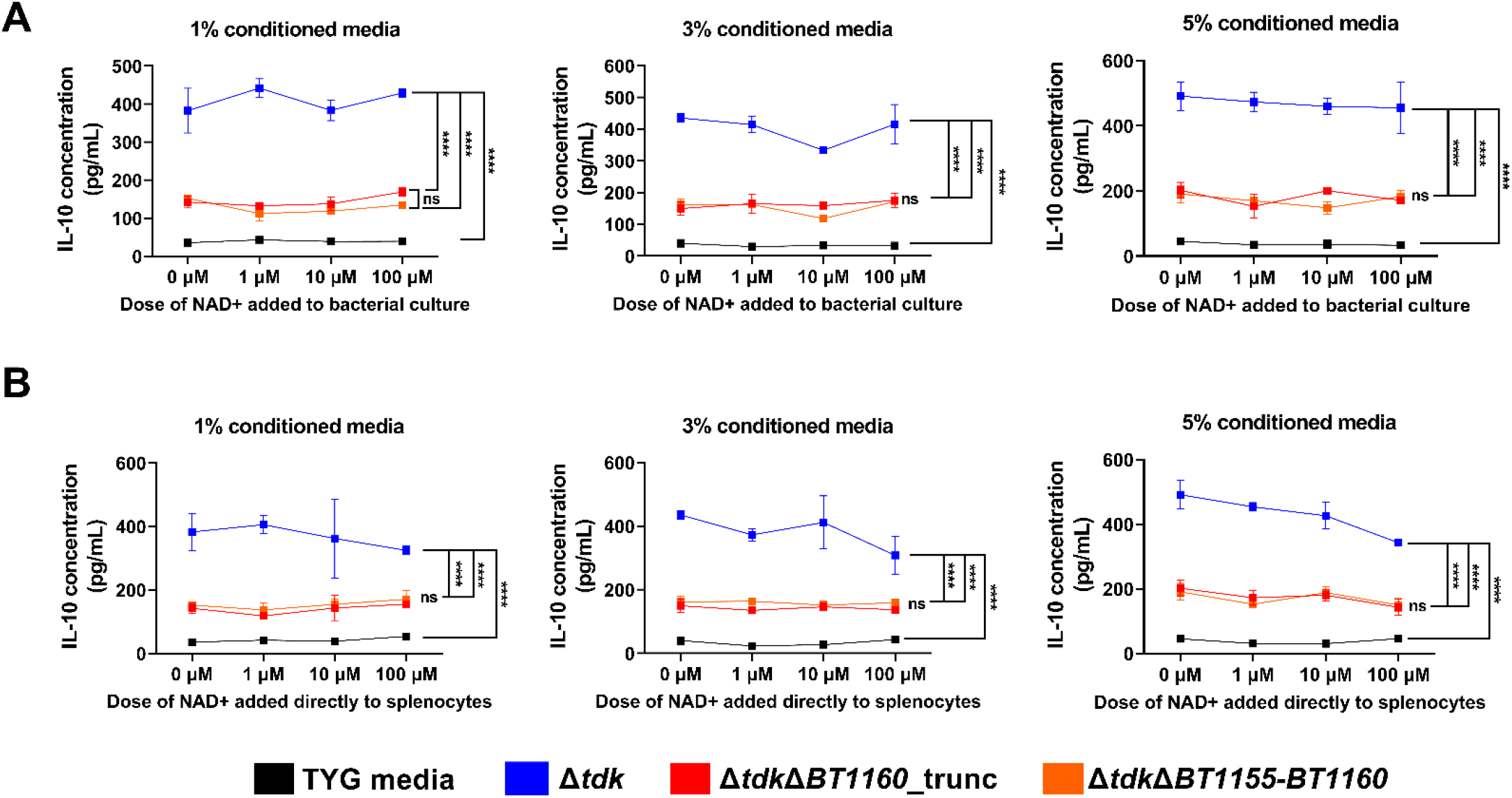
NAD+ supplementation does not restore IL-10 induction by *nqr* deficient *B. theta* mutants. **(A)** Wild-type splenocytes were stimulated with conditioned media (1%, 3%, or 5 % v/v) from wild-type *B. theta* (Δ*tdk*), *B. theta* harboring a deletion of gene *BT1160*_trunc, or genes *BT1155-BT1160*, or control TYG media for 2 days, and IL-10 levels were measured by ELISA. The conditioned media was generated from bacterial cultures that were grown in TYG media supplemented with NAD+ at a final concentration of 0 μM, 1 μM, 10 μM, or 100 μM. **(B)** Wild-type splenocytes were plated in RPMI supplemented with NAD+ to a final concentration of 0 μM, 1 μM, 10 μM, or 100 μM, and stimulated with conditioned media from wild-type *B. theta* (Δ*tdk*), *B. theta* harboring a deletion of gene *BT1160*_trunc, or genes *BT1155-BT1160*, or control TYG media (1%, 3%, or 5% v/v). Wild-type splenocytes were stimulated with for 2 days, and IL-10 levels were measured by ELISA. Data points represent the mean of two technical replicates and error bars shown the standard deviation **(A/B)**. Graphs are representative of two experiments **(A/B)**. Statistical significance was determined using two-way ANOVA with Tukey’s post hoc test and comparisons between all conditioned media groups (relevant statistical comparisons shown): **(A/B)**, ns (not significant) P≥0.05, ****P≤0.0001.

### The NQR complex influences *B. thetaiotaomicron* OMV biogenesis and composition

Since NAD+ regeneration did not seem to play a role in the ability of *B. theta* to induce IL-10, we next aimed to investigate what differed between the conditioned media from the *nqr* mutants and the parent strain (Figure 5A). As OMVs have been shown to be key mediators of the anti-inflammatory effect of other members of the *Bacteroides* (33), and a central mechanism through which *B. theta* interacts with the host (45), we reasoned that the loss of NQR could alter the production of the molecule(s) packaged into OMVs that drive IL-10 induction, or impair the production/secretion of OMVs into the conditioned media. To first determine the impact of gene *BT1160* on production of immunomodulatory OMVs we prepared OMVs from the truncated *BT1160* mutant and the parental strain, and assessed the ability of these preparations to stimulate IL-10. In these assays, OMVs were resuspended in an equivalent volume of PBS as that of the conditioned media from which they were isolated, *i.e.* if 1 mL of media was used to isolate OMVs, the pelleted vesicles were resuspended in 1 mL of PBS and not normalized based on OMV quantity. These experiments recapitulated what we observed with conditioned media, with OMV preparations from the *BT1160* deficient strain (*B. theta*^Δ*tdk*Δ*BT1160_*trunc^) showing significant impairment in IL-10 induction when compared to the parent strain (*B. theta*^Δ*tdk*^) (Figure 5B). OMVs isolated from the partial *nqr* locus deletion mutant (Δ*BT1155*-*BT1159*) that retains *BT1160* (Figure 5C) or the entire deletion mutant (Δ*BT1155-BT1160*) (Figure 5D), were also impaired in IL-10 induction ability compared to OMVs from the parent strain. However, OMVs isolated from a mutant harboring a deletion of a gene outside the *nqr* locus (Δ*BT1161*), did not alter the ability of *B. theta* OMVs to induce IL-10 compared to the parent strain (Figure 5E). This pattern was observed using male and female mice (Supplemental Figure 3B), suggesting it is not specific to a single biological sex. These data suggest that gene *BT1160* and the *nqr* locus influence the ability of *B. theta* to produce immunomodulatory OMVs.

**Figure 5.**
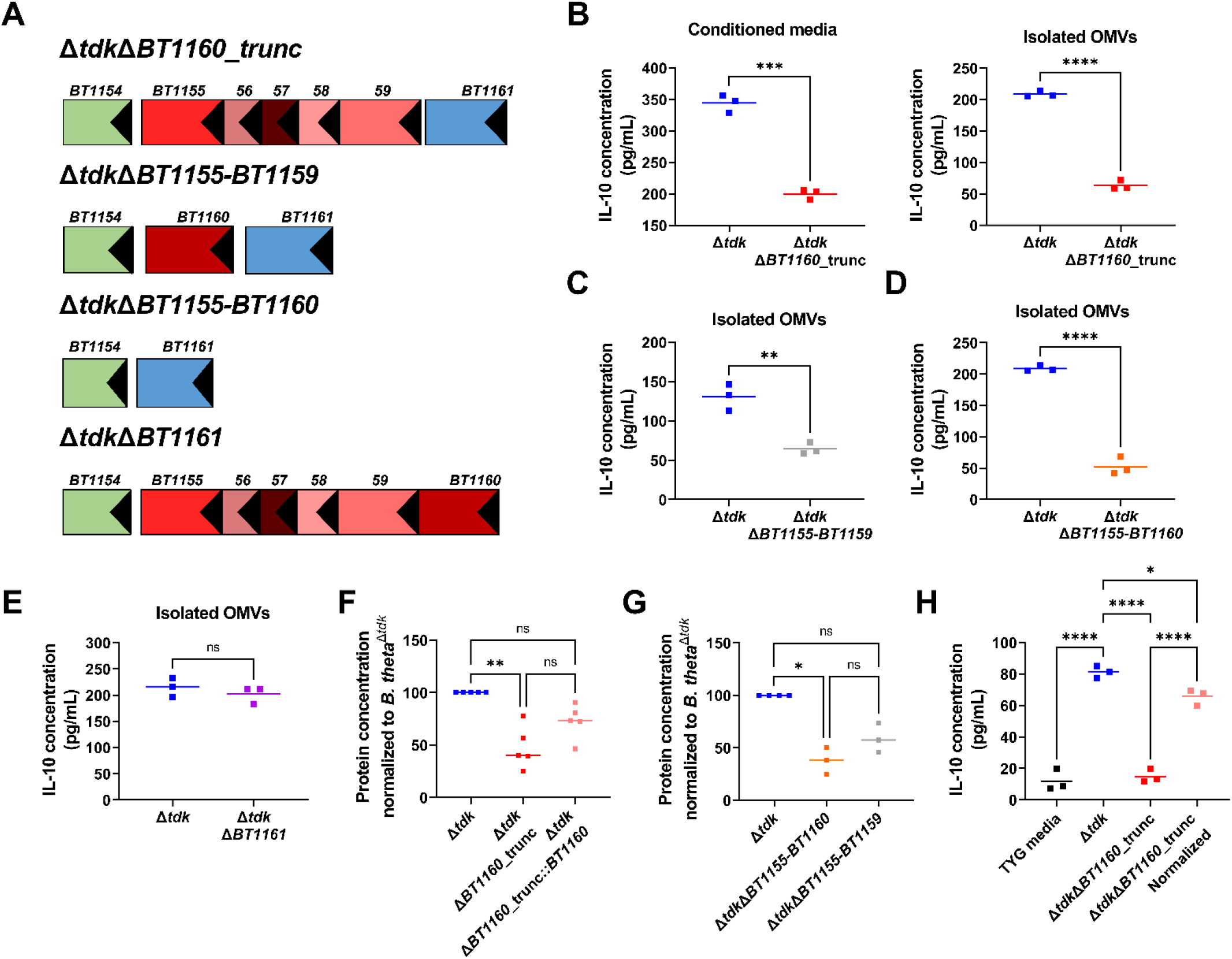
OMV production is directly linked to the ability of *B. theta* to induce IL-10 via the NQR complex. **(A)** Schematic of NQR gene locus present within each *B. theta* mutant. **(B)** Wild-type splenocytes were stimulated with conditioned media (1% v/v) or OMVs isolated from the conditioned media (1% v/v of OMVs resuspended in PBS back to original volume from which they were isolated) from *B. theta*^Δ^*^tdk^* or *B. theta*^Δ^*^tdk^*^Δ^*^BT1160_^*^trunc^ for 2 days, and IL-10 levels were measured by ELISA. **(C-E)** Wild-type splenocytes were stimulated with OMVs isolated from the conditioned media (1% v/v of OMVs resuspended in PBS as in (B)) from *B. theta*^Δ^*^tdk^* or *B. theta*^Δ^*^tdk^*^Δ^*^BT1155-BT1159^* **(C)**, *B. theta*^Δ^*^tdk^* or *B. theta*^Δ^*^tdk^*^Δ^*^BT1155-BT1160^* **(D)**, or *B. theta*^Δ^*^tdk^* or *B. theta*^Δ^*^tdk^*^Δ^*^BT1161^* **(E)** for 2 days, and IL-10 levels were measured by ELISA. **(F, G)** Protein concentration of independent biological batches of OMVs isolated from the conditioned _media of *B. theta*Δ_*_tdk_*_, *B. theta*Δ_*_tdk_*_Δ_*_BT1160__*_trunc, or *B. theta*Δ_*_tdk_*_Δ_*_BT1160__*_trunc*::BT1160* **(F)**, or *B. theta*Δ_*_tdk, B._ theta*^Δ*tdk*Δ*BT1155-BT1159*^, or *B. theta*^Δ*tdk*Δ*BT1155-BT1160*^ **(G)**, as measured by Qubit. Protein concentration was normalized to the value from a biological batch prepared concurrently from the parent strain (*B. theta*^Δ*tdk*^) which was set to 100. **(H)** Wild-type splenocytes were incubated with OMVs isolated from the conditioned media of *B. theta*^Δ^*^tdk^* or *B. theta*^Δ*tdk*Δ*BT1160_*trunc^ (1% v/v). Non-normalized OMVs were resuspended in an equivalent volume of PBS to the original volume of conditioned media they were isolated from (blue and red) and normalized OMVs were concentrated to normalize the protein concentration between the mutant and the parent strain (pink). Samples were incubated for 2 days, and IL-10 was measured in culture supernatants by ELISA. Data points represent independent technical replicates **(B-E, H)** or independent biological replicates **(F, G)** and horizontal bars represent the mean **(B-E, H)**, or median **(F, G)**. Graphs are representative of 4 experiments **(B-E, H)**, 5 pooled experiments (note, graph shows pooled data from all experiments) **(F)**, 3 pooled experiments **(G)**. Statistical significance was determined using Student’s T test: **(B-E)**, Kruskal-Wallis test with Dunn’s post-hoc test, comparisons between all: **(F, G)**, or one-way ANOVA with Dunnett’s post-hoc test, comparisons between all: **(H)**. ns (not significant) P≥0.05, **\***P<0.05, **P≤0.01, ***P≤0.001, ****P≤0.0001.

To determine if loss of *BT1160* impacted the specific proteins packaged into OMVs, including lipoproteins which are abundant in *B. theta* OMVs and known TLR2 agonists, we performed proteomic assessment of OMVs isolated from the *BT1160* truncated mutant (*B. theta*^Δ*tdk*Δ*BT1160_*trunc^) (the truncated mutant was chosen as it was a less profound modification of the *B. theta* genome yet had large effects on IL-10 induction) or the parent strain (*B. theta*^Δ*tdk*^) using LC-MS (detected proteins were identified in the reference *B. theta*^VPI-5482^ UniProtKB protein database using the program Andomeda). We performed two independent proteomic assessments with three replicates of wild-type and mutant in each analysis. A total of 187 proteins were detected in each of the two experiments (i.e. they were detected in both experiments), the majority of which (87.7%) remained unchanged between the parent and mutant strain (data not shown).The proteins that were (i) detected, (ii) found to have a fold change ≥2 in abundance, and (iii) whose P-value for differential abundance was <0.05, were considered to represent those proteins that are robustly regulated by *BT1160.* Of the 187 proteins detected, 6.42% were found to be higher in abundance in the parent strain compared to the mutant, while 5.88% were found to be higher in abundance in the *B. theta* mutant lacking *BT1160* (data not shown). Those proteins whose abundance was significantly impacted by deletion of *BT1160* from the first and second proteomics assessment are shown Supplemental Figure 3C. 14 of the 23 significantly altered proteins identified were also found to have significantly changed abundance between the OMVs isolated from the acapsular *B. theta* mutant lacking gene *BT1160* and the acapsular parent strain (data not shown). The relatively low number of altered proteins prompted us to search for alternative explanations for the impaired induction of IL-10.

Interestingly, during the preparation of OMVs for proteomic analysis, we noted that the *BT1160* deficient strain produced smaller pellets than the parent strain from equivalent amounts of conditioned media, suggesting an impairment in OMV biogenesis. Quantification of the OMVs by assessment of the protein concentration in OMV preparations revealed that deletion of gene *BT1160* leads to a reduction in the concentration of OMVs produced compared to the parent strain (Figure 5F). Note, due to variability in the raw OMV values we quantified OMVs from batches of wild-type and *B. theta* mutants that were processed at the same time, and normalized the value of the mutant to that of the wild-type which was set to 100%. The data presented show the pooled results of multiple individual biological replicate batches. Similar to the induction of IL-10 by conditioned media, complementation of gene *BT1160 in trans* restored OMV biogenesis to varying degrees (Figure 5F). As the specific method used for OMV quantitation can impact conclusions (74), we sought an alternative approach for assessment of OMV concentration. To this end we used a particle analyzer, ZetaView, to quantify OMVs. These data also showed impaired OMV production from the truncated *B. theta*^Δ*tdk*Δ*BT1160_*trunc^ mutant compared to the parent strain (Supplemental Figure 3D), validating the Qubit-based quantification data, and suggesting that the lower protein levels reflected fewer OMVs rather than a reduced protein level within the OMVs. Deletion of the remaining *nqr* locus genes except for *BT1160* (Δ*BT1155-BT1159*), and the entire *nqr* locus (Δ*BT1155-BT1160*) also led to a reduction in the concentration of OMVs compared to the parent strain (Figure 5G), suggesting that this pathway is impacted by the NQR complex rather than by gene *BT1160* in isolation. Importantly, the inability of *B. theta* lacking gene *BT1160* to induce IL-10 is not from a total loss of OMV biogenesis. ZetaView analysis and transmission electron microscopy (TEM) both revealed that *B. theta* still produces OMVs when gene *BT1160* is deleted (Supplemental Figures 3D, E), highlighting that these data reflect a quantitative reduction in OMVs rather than a complete loss of OMV production. Collectively, these data suggest that the NQR complex plays an important role in OMV biogenesis in *B. theta*.

### NQR-dependent OMV production is directly linked to the ability of *B. thetaiotaomicron* to induce IL-10

Our proteomic and OMV quantification data suggests that impaired NQR function leads to slight alterations in the specific proteins packaged in OMVs as well as a reduced capacity to generate OMVs. Thus, the impaired IL-10 induction could result from a reduction in the TLR2 agonist within the OMVs that drives the IL-10 production, a reduction in OMV quantity without changes to the specific TLR2 agonist that drives IL-10 production, or a combination of these two possibilities acting synergistically. We reasoned that if the phenotype was driven primarily by impaired OMV biogenesis, then the use of equivalent OMV concentrations from wild-type and *BT1160* mutant *B. theta* should lead to a similar amount of IL-10 induction. Alternatively, if there was impaired packaging of the specific TLR2 agonist that drives IL-10 production in OMV from the *BT1160* mutant (*B. theta*^Δ*tdk*^*^ΔBT1160_^*^trunc^), then normalizing OMV concentrations to that of the parent strain would not restore IL-10 induction by the *BT1160* mutant strain. To test this, we stimulated splenocytes with OMV preparations from the *BT1160* mutant and parental strain as above, or OMVs from the *BT1160* deletion mutant such that they were present at the same concentration as the OMVs from the parent strain. These data show that normalizing the OMV concentration between the mutant and parent strain largely restored the ability of *B. theta*^Δ*tdk*Δ*BT1160_*trunc^ to induce IL-10 (Figure 5H). This result held true whether OMVs were normalized using Qubit (protein concentration) or ZetaView (particles/mL) (data not shown). These data suggest that gene *BT1160* plays an important role in OMV biogenesis in *B. theta* and explains, in large part, how this gene—and the *nqr* locus as a whole—impacts the induction of innate immune responses by *B. theta*. Interestingly, these data also suggest that IL-10 induction by the immune system is dependent on the quantity of OMVs carrying immunomodulatory factors, and that deletion of NQR complex genes disrupts canonical OMV biogenesis. Thus, our data establishes that not only are OMVs sufficient to elicit IL-10 production, but their optimal production is required for the immunomodulatory function of *B. theta*.

## Discussion

The basis through which the immune system maintains immune tolerance has fascinated immunologists for decades, but until relatively recently, this interest has focused on the induction and maintenance of tolerance to self-antigens encoded within the host genome, or non-self as represented by organs/tissues during transplantation (6, 88, 89). However, the intestine is laden with another form of non-self, the microbiome, which also necessitates the development of immune tolerance due to the myriad benefits it provides to the host (6). A failure in immune tolerance of the microbiome can lead to development of chronic diseases like inflammatory bowel disease that severely impact host health (19, 21, 22, 27, 88–90). The intestinal immune system is therefore faced with the challenge of differentiating between potential pathogens and resident microbiome members, despite both types of microorganisms producing agonists of host-PRRs such as TLR4, TLR2, Dectin-1, etc (59, 71, 77).

While the “Danger Theory” provides a conceptual framework to understand how this is achieved, the molecular processes that underlie this decision remain enigmatic (91). Several studies have established that the gut microbiome can coordinate the induction of anti-inflammatory immune responses that limit inflammation and favor mutualism via the production of a variety of metabolites and cellular products (7, 8, 10, 12–16, 28, 30, 31, 33, 36–41, 45, 53, 77, 78, 80, 92–94). Despite this, there remains a dearth of knowledge regarding the nature of these microbiome-derived molecules and the microbial pathways that govern their production and secretion. Here, we leveraged the genetic tractability of the gut symbiont *B. theta*, a potent modulator of immune function, to identify pathways that contributed to its ability to promote the production of the anti-inflammatory cytokine IL-10. Our data revealed that secreted factors from *B. theta* promoted IL-10 production via a TLR2 and MyD88-dependent mechanism that was largely independent of TLR4. Furthermore, we uncovered that the gene, *BT1160*, and the *nqr* locus in which it resides (predicted to encode a the Na+-transporting NADH:ubiquinone oxidoreductase (NQR) complex) (55–57), played a central role in the ability of *B. theta*-derived conditioned media to drive the production of IL-10. We further show that the impairment in IL-10 induction was linked to the ability to produce outer membrane vesicles (OMVs), reinforcing the notion that OMVs play important roles as mediators of symbiosis.

The existence of a microbiome-TLR2-IL-10 axis has emerged due to the work of several groups. This is exemplified by *B.* fragilis which produces the capsular polysaccharide, polysaccharide A (PSA), which is packaged into OMVs and drives IL-10 production via TLR2 (13, 30–33, 44). Mirroring these findings, *Bifidobacterium bifidum* (14) and *Helicobacter hepaticus* (15) are purported to produce cellular polysaccharides that stimulate IL-10 production via TLR2, and members of the Bacteroidetes as a whole have recently been shown to promote IL-10 induction through a TLR2-MyD88 axis (80). Our data reveal that *B. theta* secreted factors also stimulated IL-10 production through a TLR2 and MyD88-dependent mechanism. Notably, it has recently emerged that chemically synthesized PSA lacks IL-10 inductive capacity, by contrast with PSA isolated from *B. fragilis*, and instead, a small lipid covalently attached to PSA is critical for its immunomodulatory function (30), in accordance with structural studies of TLR2-based recognition of synthetic agonists that show that TLR2 directly binds lipids (70, 95). We did not detect a role for Dectin-1 in promoting IL-10 production, which contrasts with *B. fragilis* whereby dual engagement of TLR2 by lipid and Dectin-1 by PSA stimulate IL-10 (30), suggesting that *B. theta* shapes immune function by a pathway distinct from *B. fragilis*. Notably, neither *H. hepaticus* nor *B. bifidum* require sensing by Dectin-1 for their immunomodulatory capacity (14, 15). It is therefore unclear if there are as yet unidentified lipids/lipid moieties that are required for the immunomodulatory function of *H. hepaticus* or *B. bifidum* derived polysaccharides, and further investigation is required to identify the specific factor from *B. theta* that stimulates IL-10 induction. Given the lack of molecularly defined factors that drive IL-10, it is as yet unknown if TLR2 represents a common host sensor for a wide variety of microbiome members through recognition of distinct molecules, or if there is a common structurally similar lipid produced across a range of microbes that promotes IL-10 induction. Moreover, whether specific chemical features of gut microbe-derived molecules are sensed to allow induction of tolerance, or other facets of the initial recognition event are responsible for the observed responses is unknown and warrants intensive study.

A number of studies have shown OMVs to be sufficient to mediate the immunologic function of gut bacteria (33, 35, 45, 48, 53, 96). Our data now provide evidence that OMV production is directly linked to immunomodulation and that impaired OMV production can reduce the capacity to shape immune function. Our data identified the existence of a complex, namely that encoded by *nqr*, which is required for optimal production of OMVs, and demonstrated that this OMV biogenesis could be directly linked to immunomodulatory capacity. Despite the attention that has been focused on OMVs (49, 97, 98), there is still a dearth of information regarding the molecular pathways that coordinate their biogenesis. Seminal studies on *B. theta* OMVs have revealed that they are composed of phospholipids, lipooligosaccharide (LOS), outer membrane and periplasmic proteins, sphingolipids, and lipoproteins (96, 99). These OMVs contain distinct protein profiles from the *B. theta* outer membrane and differ from other microbes in that they are highly enriched with lipoproteins (96, 100–102). The OMV-enriched lipoproteins contain a lipoprotein export sequence believed to coordinate packaging into OMVs, suggesting the existence of an ordered process (96). Our studies extend on these and identify a locus that sustains the optimal production of OMVs. The *nqr* locus supports energy production during respiration, and in doing so, shuttles Na+ across the membrane to create Na+ gradients, and regenerates NAD+ (an electron acceptor) from NADH (55–57). While we currently do not understand which of these functions, if any, are involved in OMV biogenesis, our studies show that regeneration of NAD+ is likely not the predominant mechanism. It is therefore tempting to speculate that energetic requirements of OMV generation necessitate optimal ATP generation, and hence a functional NQR complex is essential.

Collectively, our studies have identified a novel regulator of the immunomodulatory function of *B. theta*. The capacity to modulate OMV biogenesis has applicability to both promote tolerance and limit pathogenicity by increasing or decreasing OMV production in mutualists or pathogens respectively, but this will require detailed understanding of how NQR contributes to this process. Notably, antimicrobial compounds such as natural quinone-like inhibitors—HQNO and korormicin—target the NQR complex by binding to NqrB and preventing Na+-dependent reduction of quinones (103). These compounds effectively target and kill pathogenic species, like *Vibrio cholerae* and *Pseudomonas aeruginosa*, yet gut commensals like *B. fragilis* and *B. theta* remain unharmed (103). Thus, the selective inhibition or stimulation of the NQR complex may allow rational manipulation of microbiome function to enhance anti-inflammatory function, or boost the delivery of molecules of interest via OMVs. Further insight into how OMV biogenesis is coordinated, and defining the specific immunomodulatory factors they harbor is essential to gain a more complete understanding of how gut microbes like *B. theta* coordinate host immune responses, and will provide an opportunity to identify pathways that can be manipulated to shape intestinal immune responses.

## Acknowledgments

We are grateful to Anagha Kadam, Orion Brock, Adeline Hajjar, and Jan Claesen for their intellectual input throughout these studies, and their advice during preparation of the manuscript; Meng Wu for her assistance with data analysis from the transposon mutagenesis screen; Belinda Willard for proteomic analysis; and William Massey for his assistance with generation of bone-marrow derived macrophages. In addition, we would like to thank the following for their provision of mice used in these studies: Danielle Kish and Robert Fairchild, Cleveland Clinic Lerner Research Institute (TLR4^-/-^), Nancy Nagy, Case Western Reserve University (TLR2^-/-^, MyD88^-/-^), and Luis Ramirez and Tanya Freedman, University of Minnesota (wild-type and Dectin-1^-/-^). OMV proteomics was made possible by the Fusion Lumos LC-MS instrument purchased with a National Institutes of Health (NIH) Shared Instrument Grant: 1S10OD023436-01.

## Author Contributions

M.J.E. designed and executed experiments to test the role of various forms of *B. theta* to induce IL-10, identified transposon mutants with impaired IL-10 induction capacity, generated mutant forms of *B. theta*, analyzed data and helped write the manuscript. R.W.P.G. generated and grew the transposon mutagenesis library used in this study, generated mutant forms of *B. theta*, and designed experiments to test the role of the *nqr* locus. J.M.T assisted in executing experiments and provided intellectual input for data interpretation. C.V.H. provided TLR2^-/-^ and MyD88^-/-^ mice and guidance on their use in these studies. E.C.M. supervised the development of the transposon mutagenesis library, and provided the library and acpasular mutant of *B. theta* for use in our studies, in addition to intellectual input for data interpretations. P.P.A. designed experiments described within, supervised the study, and helped write the manuscript.

