## Supplemental_Figures for "The NQR complex regulates the immunomodulatory function of *Bacteroides thetaiotaomicron*"

### Supplemental Figure 1.

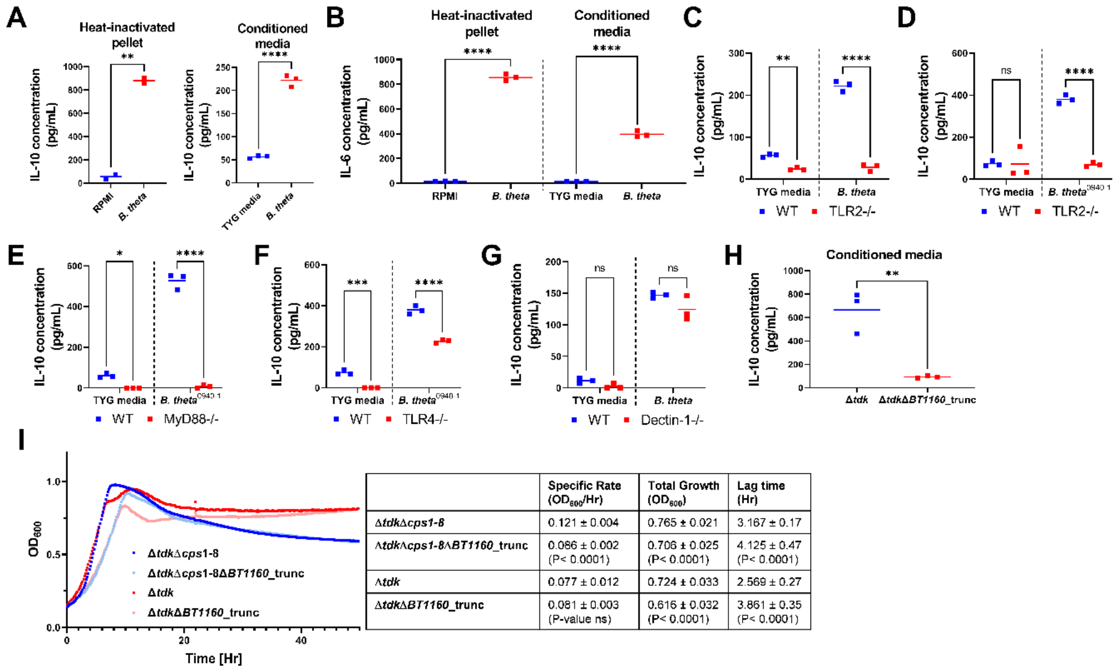


| | Specific Rate ($OD_{600}$/Hr) | Total Growth ($OD_{600}$) | Lag time (Hr) |
|---|---|---|---|
| *Δtdk*Δ*cps1-8* | 0.121 ± 0.004 | 0.765 ± 0.021 | 3.167 ± 0.17 |
| *Δtdk*Δ*cps1-8*Δ*BT1160*_trunc | 0.086 ± 0.002 (P< 0.0001) | 0.706 ± 0.025 (P< 0.0001) | 4.125 ± 0.47 (P< 0.0001) |
| *Δtdk* | 0.077 ± 0.012 | 0.724 ± 0.033 | 2.569 ± 0.27 |
| *Δtdk*Δ*BT1160*_trunc | 0.081 ± 0.003 (P-value ns) | 0.616 ± 0.032 (P< 0.0001) | 3.861 ± 0.35 (P< 0.0001) |

**Supplemental Figure 1. Cytokine induction by *B. theta* strains or *BT1160* mutants on wild-type, TLR2$^{-/-}$, MyD88$^{-/-}$, TLR4$^{-/-}$, and Dectin-1$^{-/-}$ splenocytes and BMDMs, and *BT1160* mutant (on acapsular or wild-type backgrounds) and parent strain growth curves, related to Figures 1 and 2.**

**(A)** Wild-type BMDMs were stimulated with *B. theta*$^{VPI-5482}$ heat-inactivated pellet ($OD_{600}$=1.0) or RPMI media, or *B. theta*$^{VPI-5482}$ conditioned media or TYG media (1% v/v) control for 2 days and IL-10 was assessed in the supernatant by ELISA.
**(B)** Wild-type splenocytes were stimulated with *B. theta*$^{VPI-5482}$ heat-inactivated pellet ($OD_{600}$=1.0) or RPMI media, or *B. theta*$^{VPI-5482}$ conditioned media or TYG media control (1% v/v) for 2 days and IL-6 was assessed in the supernatant by ELISA.
**(C)** Wild-type and TLR2$^{-/-}$ BMDMs were stimulated with *B. theta*$^{VPI-5482}$ conditioned media or TYG media control (1% v/v) for 2 days and IL-10 was assessed in the supernatant by ELISA.
**(D-F)** Wild-type and TLR2$^{-/-}$ **(D)**, MyD88$^{-/-}$ **(E)**, or TLR4$^{-/-}$ **(F)** splenocytes were stimulated with conditioned media (1% v/v) from *B. theta* (strain 0940-1) or TYG media control for 2 days and IL-10 was assessed in the supernatant by ELISA.
**(G)** Wild-type and Dectin-1$^{-/-}$ splenocytes were stimulated with *B. theta*$^{VPI-5482}$ conditioned media (1% v/v) or TYG media control for 2 days and IL-10 was assessed in the supernatant by ELISA.
**(H)** Wild-type BMDMs were stimulated with conditioned media (1% v/v) from *B. theta* wild-type (Δ*tdk*) and the *B. theta* (Δ*tdk*Δ*BT1160*_trunc) for 2 days and IL-10 was assessed in the supernatant by ELISA.
**(I)** *B. theta* wild-type (Δ*tdk*) and acapsular *B. theta* (Δ*tdk*Δ*cps1-8*) and *B. theta* with the truncated deletion of gene *BT1160* on each background were grown in TYG for growth curve analysis. Growth curves were performed in 96 well plates with 12 technical replicates per mutant. Growth ($OD_{600}$) was measured continuously over 48 hours with readings every 10 minutes. Table represents the average and standard deviation of the Specific Rate, Total growth, and Lag Time of all 12 technical replicates for each mutant. Statistical significance calculated comparing the mutant to the appropriate parent strain and reported in the tables under the mutant.

Data points represent an independent technical replicate where the horizontal bars represent the mean **(A-H)**, or the average of 12 technical replicates **(I)**. Graph is representative of at least 2 experiments **(A-I)**. Statistical significance was determined Student's T test: **(A, B, H, I)** or two-way ANOVA with Sidak's post hock test, comparisons to wild-type **(C-G)**. ns P≥0.05, *P<0.05, **P≤0.01, ***P≤0.001****P≤0.0001.

**Supplemental Figure 2.**

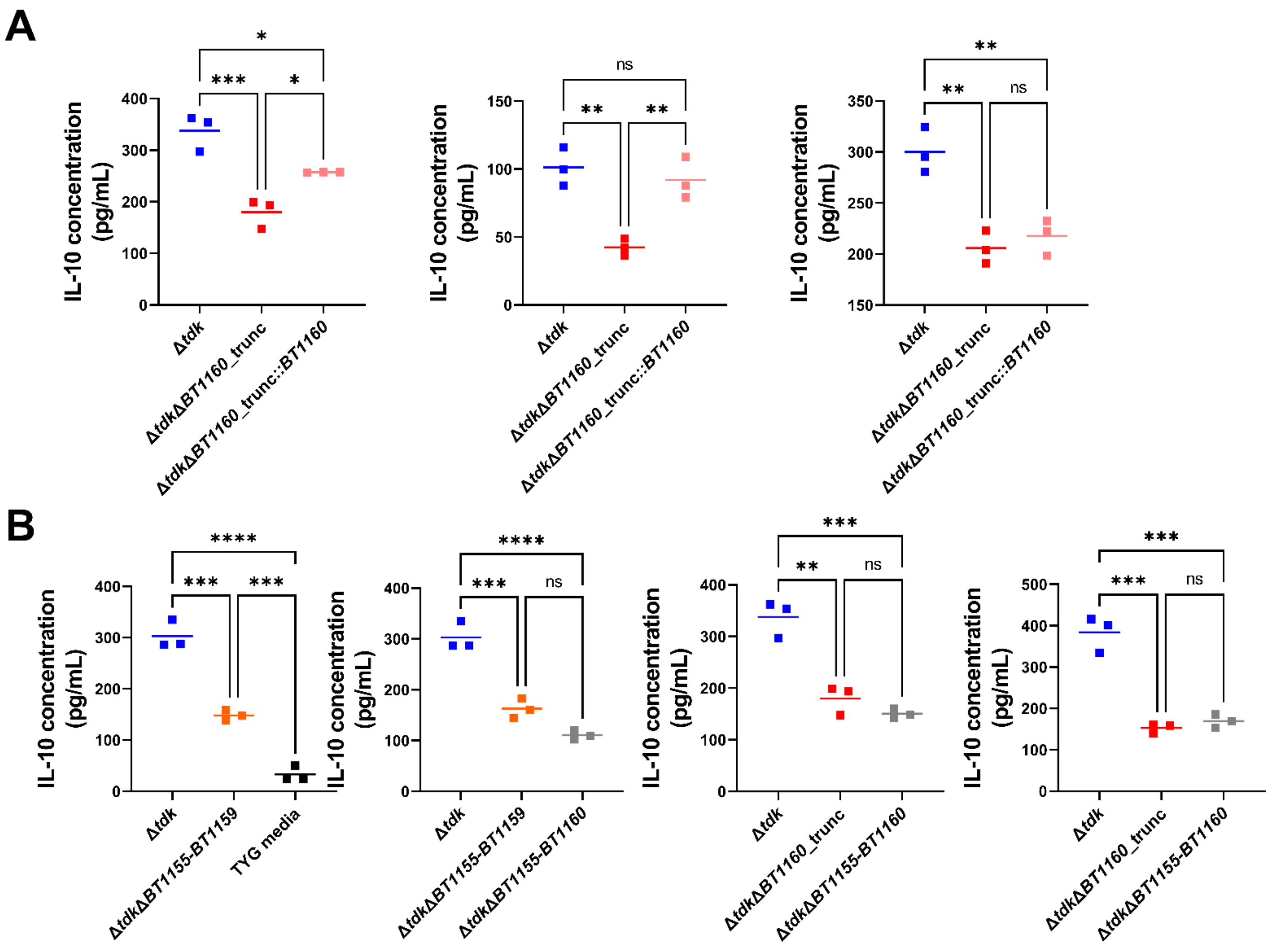


**Supplemental Figure 2. Additional biological replicates of *B. theta BT1160* complementation and *nqr* locus mutants, related to Figures 2 and 3.**

**(A)** Wild-type splenocytes were stimulated with *B. theta*$^{\Delta tdk}$, *B. theta*$^{\Delta tdk \Delta BT1160\_\mathrm{trunc}}$, or *B. theta*$^{\Delta tdk \Delta BT1160\_\mathrm{trunc}::BT1160}$ conditioned media (1% v/v) for 2 days and IL-10 was assessed in the supernatant by ELISA. Each graph shows the results from an independent experiment.

**(B)** Wild-type splenocytes were stimulated with *B. theta*$^{\Delta tdk}$, *B. theta*$^{\Delta tdk \Delta BT1160\_\mathrm{trunc}}$, *B. theta*$^{\Delta tdk \Delta BT1155\text{-}BT1159}$, or *B. theta*$^{\Delta tdk \Delta BT1155\text{-}BT1160}$ conditioned media or TYG media control (1% v/v) for 2 days and IL-10 was assessed in the supernatant by ELISA. Each graph shows the results from an independent experiment.

Data points on each graph represent an independent technical replicate where the horizontal bars represent the mean **(A, B)**. Statistical significance was determined using one-way ANOVA with Dunnett's post-hoc test, comparisons between all: **(A, B)**. ns P≥0.05, ***P<0.05, **P≤0.01, ***P≤0.001, ****P≤0.0001.

**Supplemental Figure 3.**

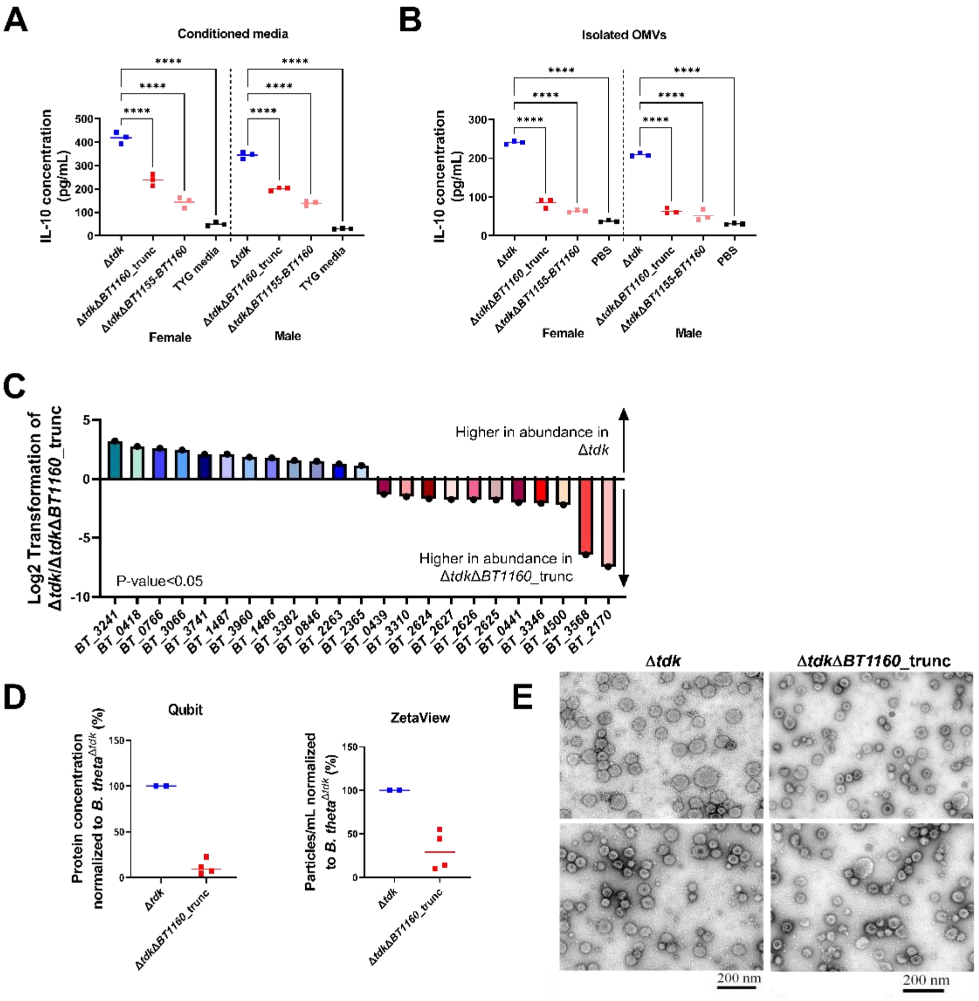


**Supplemental Figure 3. IL-10 induction by *B. theta nqr* locus mutants on wild-type splenocytes isolated from male and female mice, and OMV proteomics, concentration measurement, and images, related to Figure 3 and 5.**

**(A, B)** Wild-type splenocytes, isolated from male and female mice, were stimulated with *B. theta*$^{\Delta tdk}$, *B. theta*$^{\Delta tdk\Delta BT1160\_trunc}$, or *B. theta*$^{\Delta tdk\Delta BT1155\text{-}BT1160}$ conditioned media or TYG control media (1% v/v) **(A)** or isolated OMVs (1% v/v; resuspended in PBS) **(B)** for 2 days and IL-10 was assessed in the supernatant by ELISA.
**(C)** Proteins with significant (P-value ≤0.05) differential abundance and with fold changes ≥2, identified in two separate OMV proteomics analyses between *B. theta*$^{\Delta tdk}$ and *B. theta*$^{\Delta tdk\Delta BT1160\_trunc}$ are shown. Proteins identified are represented by the gene locus tag. Fold change abundance is representative of one experiment. Bars represents the mean difference in abundance in OMVs from *B. theta*$^{\Delta tdk}$ relative to *B. theta*$^{\Delta tdk\Delta BT1160\_trunc}$.
**(D)** Qubit and ZetaView were used to measure the concentration of OMVs isolated from the same batch of conditioned media of *B. theta*$^{\Delta tdk}$ and *B. theta*$^{\Delta tdk\Delta BT1160\_trunc}$. Each data point represents an independent biological replicate where the horizontal bars represent the median.
**(E)** Representative TEM images of concentrated OMVs isolated from the conditioned media of *B. theta*$^{\Delta tdk}$ and *B. theta*$^{\Delta tdk\Delta BT1160\_trunc}$.

Data points represent an independent technical replicate where the horizontal bars represent the mean **(A, B)**. Statistical significance was determined using one-way ANOVA with Dunnett's post-hoc test, comparisons to *B. theta*$^{\Delta tdk}$: **(A, B)**. ****P≤0.0001.
